# Protection from SARS-CoV-2 Delta one year after mRNA-1273 vaccination in nonhuman primates is coincident with an anamnestic antibody response in the lower airway

**DOI:** 10.1101/2021.10.23.465542

**Authors:** Matthew Gagne, Kizzmekia S. Corbett, Barbara J. Flynn, Kathryn E. Foulds, Danielle A. Wagner, Shayne F. Andrew, John-Paul M. Todd, Christopher Cole Honeycutt, Lauren McCormick, Saule T. Nurmukhambetova, Meredith E. Davis-Gardner, Laurent Pessaint, Kevin W. Bock, Bianca M. Nagata, Mahnaz Minai, Anne P. Werner, Juan I. Moliva, Courtney Tucker, Cynthia G. Lorang, Bingchun Zhao, Elizabeth McCarthy, Anthony Cook, Alan Dodson, Prakriti Mudvari, Jesmine Roberts-Torres, Farida Laboune, Lingshu Wang, Adrienne Goode, Swagata Kar, Seyhan Boyoglu-Barnum, Eun Sung Yang, Wei Shi, Aurélie Ploquin, Nicole Doria-Rose, Andrea Carfi, John R. Mascola, Eli A. Boritz, Darin K. Edwards, Hanne Andersen, Mark G. Lewis, Mehul S. Suthar, Barney S. Graham, Mario Roederer, Ian N. Moore, Martha C. Nason, Nancy J. Sullivan, Daniel C. Douek, Robert A. Seder

## Abstract

mRNA-1273 vaccine efficacy against SARS-CoV-2 Delta wanes over time; however, there are limited data on the impact of durability of immune responses on protection. We immunized rhesus macaques at weeks 0 and 4 and assessed immune responses over one year in blood, upper and lower airways. Serum neutralizing titers to Delta were 280 and 34 reciprocal ID_50_ at weeks 6 (peak) and 48 (challenge), respectively. Antibody binding titers also decreased in bronchoalveolar lavage (BAL). Four days after challenge, virus was unculturable in BAL and subgenomic RNA declined ∼3-log_10_ compared to control animals. In nasal swabs, sgRNA declined 1-log_10_ and virus remained culturable. Anamnestic antibody responses (590-fold increase) but not T cell responses were detected in BAL by day 4 post-challenge. mRNA-1273-mediated protection in the lungs is durable but delayed and potentially dependent on anamnestic antibody responses. Rapid and sustained protection in upper and lower airways may eventually require a boost.

## INTRODUCTION

COVID-19 vaccines designed to express the spike (S) protein of SARS-CoV-2, including the mRNA-based vaccines mRNA-1273 (Baden et al., 2021a) and BNT162b2 (Polack et al., 2020) and the adenovirus-vectored vaccines Ad26.COV2.S (Sadoff et al., 2021) and AZD1222 (Ramasamy et al., 2021), have shown substantial protection against vaccine-matched virus strains. mRNA-1273 had an efficacy of 94% in a Phase III clinical trial (Baden et al., 2021a) and 96.3% among US healthcare workers (Pilishvili et al., 2021). mRNA-1273-elicited neutralizing antibodies were still detectable six months after immunization (Doria-Rose et al., 2021). However, new SARS-CoV-2 variants have mutations that decrease the sensitivity of vaccine-elicited neutralization and increase viral replication, raising concerns about the durability of protection provided by mRNA-1273 and other vaccines (Baden et al., 2021b; Bruxvoort et al., 2021; Puranik et al., 2021).

Delta (henceforth referred to by its Pango lineage, B.1.617.2) is currently the dominant circulating strain of SARS-CoV-2 and a WHO-designated variant of concern (VOC). This strain was first identified in India in October 2020 amidst substantial levels of community transmission (Cherian et al., 2021; Mishra et al., 2021; Mlcochova et al., 2021). B.1.617.2 contains the mutations L452R, T478K, D614G, and P681R in the receptor-binding domain (RBD) and C-terminus of the S1 binding subdomain. These substitutions contribute to both increased receptor binding and reduced neutralization by vaccine-elicited and monoclonal antibodies (mAbs) (Ozono et al., 2021; Planas et al., 2021; Tada et al., 2021; Wang et al., 2021b). In addition, B.1.617.2 has acquired several unique mutations in the N-terminal domain (NTD) that substantially decrease antibody binding and neutralization sensitivity including T19R, G142D, and a deletion at positions 156-158 accompanied by a G insertion (Liu et al., 2021; Planas et al., 2021). Neutralizing antibody titers from mRNA-1273 and BNT162b2 vaccinee sera to B.1.617.2 are reduced 3-fold compared to the vaccine-matched strain, USA-WA1/2020 (WA1) shortly after immunization (Edara et al., 2021b). A 7-fold reduction in neutralizing titers for Ad26.CoV2.S sera in comparison to WA1 or WA1 with a D614G substitution (henceforth referred to as D614G) has also been described (Barouch et al., 2021; Tada et al., 2021).

Recent studies in the United Kingdom, United States, and Qatar have shown reduced efficacy of mRNA-based vaccines against asymptomatic and symptomatic, but not severe, B.1.617.2 infection (Bruxvoort et al., 2021; Chemaitelly et al., 2021; Lopez Bernal et al., 2021; Puranik et al., 2021; Tang et al., 2021). We and others have found that binding and neutralizing antibody titers in NHP and humans are key correlates of protection for mRNA and adenovirus-vectored COVID-19 vaccines (Corbett et al., 2021b; Gilbert et al., 2021; Khoury et al., 2021; Roozendaal et al., 2021). Antibody titers significantly decrease over a 6-month interval after the initial immunization series (Canaday et al., 2021; Corbett et al., 2021a; Levin et al., 2021). Retrospective analysis in Israel found that breakthrough cases in BNT162b2 vaccinees during a period of substantial B.1.617.2 transmission were statistically correlated with the length of time elapsed since vaccination, suggesting a role for waning antibody titers in vaccine efficacy reduction (Goldberg et al., 2021). Similarly, participants in the mRNA-1273 efficacy trial (COVE) who initially received a placebo prior to vaccination had reduced rates of B.1.617.2 breakthrough infections and severe disease compared to study participants who received mRNA-1273 at an earlier time (Baden et al., 2021b). These observations were consistent with previous findings from Israel showing that breakthrough infections with Alpha (B.1.1.7) were associated with lower BNT162b2-elicited binding and neutralizing antibody titers immediately prior to infection (Bergwerk et al., 2021). While BNT162b2 efficacy against breakthrough infections with B.1.617.2 has been estimated as 42% in Minnesota, USA and 53.5% in Qatar, those same studies reported mRNA-1273 efficacy as 76% and 84.8%, respectively (Puranik et al., 2021; Tang et al., 2021). Likewise, additional analysis in California, USA indicates a mRNA-1273 vaccine efficacy against B.1.617.2 of 87% (Bruxvoort et al., 2021). While any differences between mRNA-1273 and BNT162b2 may diminish with increased time since vaccination, this observation warrants continued investigation. Furthermore, there is no analysis of mRNA-1273-elicited immunity out to one year in the context of protection against mild and severe disease in the upper and lower airways.

The nonhuman primate (NHP) model has been extensively used to assess a variety of vaccine candidates against SARS-CoV-2 (Corbett et al., 2020; Francica et al., 2021; He et al., 2021; Mercado et al., 2020; van Doremalen et al., 2020) and has been reliable for predicting protective efficacy with mRNA-1273 in humans (Corbett et al., 2021b; Gilbert et al., 2021). Thus, this model is an ideal tool for examining the effect of waning antibody titers on long term protection in the context of a challenge with B.1.617.2.

Here, we immunized rhesus macaques with 100µg mRNA-1273 at weeks 0 and 4 and then challenged them with B.1.617.2 approximately one year later. To provide insights into potential mechanisms of protection, we assessed B.1.617.2-binding antibody titers from the blood and both the upper and lower airways after both vaccination and challenge. Serum neutralizing titers, and longevity of virus-specific memory B and T cell responses were also analyzed.

## RESULTS

### mRNA-1273 immunization elicits binding and neutralizing antibodies to B.1.617.2

Indian-origin rhesus macaques (n = 8/group) were immunized with 100µg mRNA-1273 or mRNA control at weeks 0 and 4 (Fig. S1A). Serum IgG binding titers to prefusion stabilized S protein (S-2P) of the vaccine-matched WA1 strain were assessed at weeks 6 (peak), 24 (memory) and 48 weeks (memory and time of challenge) after vaccination. Geometric mean titers (GMT) significantly decreased from 3000 WHO International Standard binding antibody units (BAU)/mL at week 6 to 260 BAU/mL at week 24. However, the rate of decline was less for the remainder of the year, reaching 188 BAU/mL at week 48 (Fig. 1A). 7/8 animals in the mRNA-1273 group had higher binding titers than all control NHP at week 48. Similarly, the kinetics observed for WA1 RBD-binding titers showed a 130-fold reduction in geometric mean area under curve (AUC) between weeks 6 and 25, but only an additional 3-fold reduction by week 48 (Fig. 1B). Compared to WA1 RBD, GMT at week 6 were reduced 5.4-fold to B.1.617.2 RBD (Fig. 1C). Despite the lower binding titers to B.1.617.2, antibodies were clearly functional, as sera from vaccinated NHP inhibited almost 100% of binding between the SARS-CoV-2 receptor, angiotensin-converting enzyme 2 (ACE2), and S-2P of both variants at week 6 (Fig. S1B-C).

**Figure 1.**
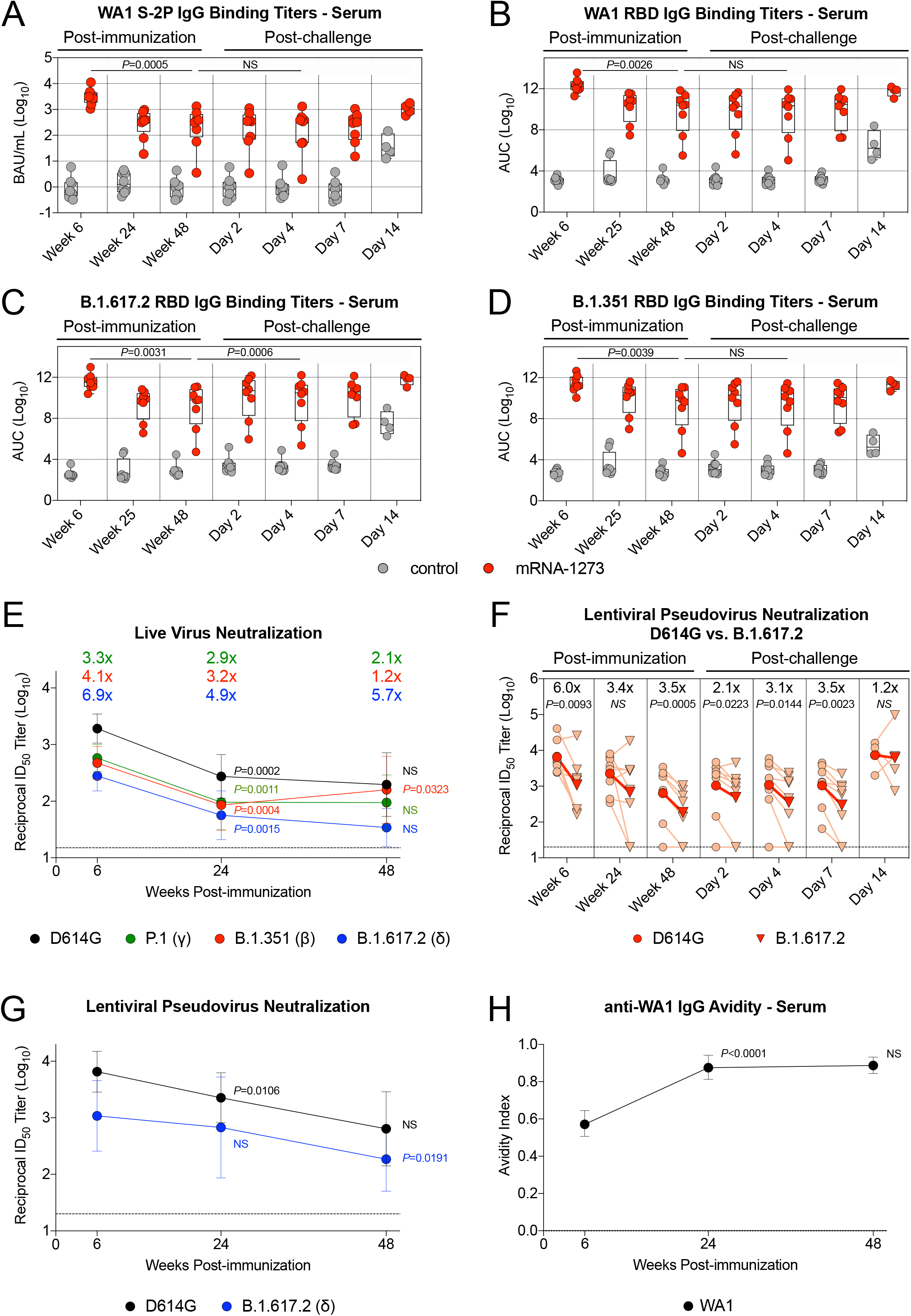
mRNA-1273 elicits SARS-CoV-2 binding and neutralizing antibodies. (A-D) Sera were collected at weeks 6, 24 or 25, and 48 post-immunization, and days 2, 4, 7 and 14 post-challenge. IgG binding titers to (A) WA1 S-2P, (B) WA1 RBD, (C) B.1.617.2 RBD, and (D) B.1.351 RBD. IgG binding titers in (A) are expressed in WHO units. Circles in (A-D) indicate individual NHP. Boxes represent interquartile range with the median denoted by a horizontal line. 4-8 NHP per group. Dotted lines in (A-D) are for visualization purposes and denote 1-log_10_ (A) or 4-log_10_ (B-D) increases in binding titers. Statistical analysis shown for mRNA-1273 cohort only. (E) Sera were collected at weeks 6, 24 and 48 post-immunization. Neutralizing antibody titers assessed from a live virus assay with D614G, P.1, B.1.351, and B.1.617.2. Circles indicate GMT, and error bars represent 95% CI. Values above circles represent fold reduction in GMT for each variant in comparison to D614G at each timepoint. Dotted line indicates assay limit of detection. 8 NHP per group. Statistical analysis shown for variant-specific responses at week 24 and 48 in comparison to corresponding variant-specific responses at preceding timepoint. (F-G) Sera were collected at weeks 6, 24 and 48 post-immunization, and days 2, 4, 7 and 14 post-challenge. Neutralizing antibody titers assessed from a lentiviral pseudovirus assay with either D614G or B.1.617.2 S. (F) Light red lines and symbols indicate individual NHP while dark red lines and symbols indicate GMT. Values above symbols represent fold reduction in neutralizing titers to B.1.617.2 compared to D614G at each timepoint. Dotted line indicates assay limit of detection. Statistical analysis shown for neutralizing titers to B.1.617.2 in comparison to D614G at each timepoint. (G) Circles indicate GMT, and error bars represent 95% CI. Dotted line indicates assay limit of detection. Statistical analysis shown for variant-specific responses at week 24 and 48 in comparison to corresponding variant-specific responses at preceding timepoint. 4-8 NHP per group. (H) Sera were collected at weeks 6, 24 and 48 post-immunization to determine anti-WA1 S-2P IgG avidity index. Circles indicate GMT, and error bars represent 95% CI. Statistical analysis shown for avidity at week 24 and 48 in comparison to avidity at preceding timepoint. 8 NHP per group. See also Table S1 for list of mutations in MULTI-ARRAY ELISA and Figure S1 for live virus neutralizing responses from individual NHP and experimental schematic.

We next measured binding titers to Beta (B.1.351) as this variant has the greatest impact on reducing neutralization by vaccinee and convalescent sera (Geers et al., 2021; Tada et al., 2021; Wang et al., 2021c), which may have contributed to reduced vaccine efficacy in regions with substantial circulation of B.1.351 (Madhi et al., 2021; Shinde et al., 2021). There was an 8-fold reduction in binding titers to B.1.351 RBD at week 6 as compared to WA1 RBD (Fig. 1D). At week 48, GMT for both B.1.351 and B.1.617.2 were similar and only 3.9-fold less than for WA1.

Serum neutralization of D614G and a panel of variants was measured using a live virus assay. mRNA-1273-vaccinated NHP had GMT to D614G of 1900 at week 6, which declined to 275 at week 24 and 200 at week 48 (Fig. 1E). The kinetics of a rapid decline in neutralizing antibody titers during the first 6 months (*P*=0.0002) followed by a slower decline at approximately 1 year (*P*>0.05) was consistent with the trend seen in binding titers (Fig. 1A-D). Compared to D614G, neutralizing GMT to Gamma (P.1), B.1.351 and B.1.617.2 were reduced 3.3-fold, 4.1-fold and 6.9-fold, respectively, at week 6. Neutralizing titers to B.1.617.2 declined over the following year, with 3/8 NHP having undetectable responses against B.1.617.2 at week 48 (Fig. S1F). Of note, neutralizing titers to B.1.351 showed a modest increase from week 24 to 48 (*P*=0.0323). This difference between the kinetics of neutralizing titers to B.1.617.2 and B.1.351 suggests a change in serum epitope dominance and/or increased antibody affinity maturation, both of which would support recent findings on the continued evolution of antibody responses induced by mRNA and adenovirus-vectored vaccines (Barouch et al., 2021; Corbett et al., 2021a; Lopez Ledesma et al., 2021).

The decrease in serum neutralization capacity over time was confirmed with a lentiviral pseudovirus assay. At week 6, pseudovirus neutralizing GMT to B.1.617.2 were reduced 6-fold as compared to D614G (*P*=0.0093) and detectable in all NHP. At weeks 24 and 48, the GMT reduction declined to 3.5-fold, and 2/8 vaccinated NHP had undetectable neutralizing titers to B.1.617.2 (Fig. 1F-G).

Finally, to further assess qualitative changes in antibody responses over time, antibody avidity was measured from sera over 48 weeks (Fig. 1H). The geometric mean avidity index of WA1 S-2P-binding IgG antibodies increased from 0.6 at week 6 to 0.9 at week 24 (*P*<0.0001). Antibody avidity remained unchanged from week 24 to week 48 (*P*>0.05).

### Serum epitope analysis reveals immuno-focusing on epitopes associated with neutralization of B.1.617.2

To extend the qualitative analysis of antibody neutralization and identify antibodies that could be contributing to avidity, we performed serum antibody epitope mapping at weeks 6, 24, and 48 after immunization. Using a surface plasmon resonance-based (SPR) competition assay, we determined the relative proportion (percent competition) of serum antibodies targeting cross-reactive epitopes on both WA1 and B.1.617.2 SARS-CoV-2 S-2P (Table S2).

Cross-reactive antigenic sites A, B, C, E, F, and G were defined by mAbs B1-182, CB6, A20-29.1, LY-COV555, A19-61.1, and S309 respectively, all targeting the RBD of SARS-CoV-2 S, with sites A, B, and F being associated with neutralization of SARS-CoV-2 B.1.617.2 (Table S2) (Corbett et al., 2021a; Wang et al., 2021b). Analysis of relative serum reactivity 48 weeks after immunization showed significantly higher proportions of serum antibodies targeting antigenic sites G, B, and F (represented by mAbs S309, CB6, and A19-61.1, respectively) on B.1.617.2 S-2P, as compared to WA1 (G: *P*=0.004; B: *P*=0.0009; F: *P*<0.0001) (Fig 2A). Longitudinal analysis of the epitope-specific serum antibody responses to WA1 S-2P showed no significant differences in the relative serum reactivity over time (Fig 2B). This indicates that while quantity of serum antibodies targeting WA1 S-2P decreased over time (Fig 1A), the composition of serum antibodies targeting cross-reactive epitopes remained unchanged.

**Figure 2.**
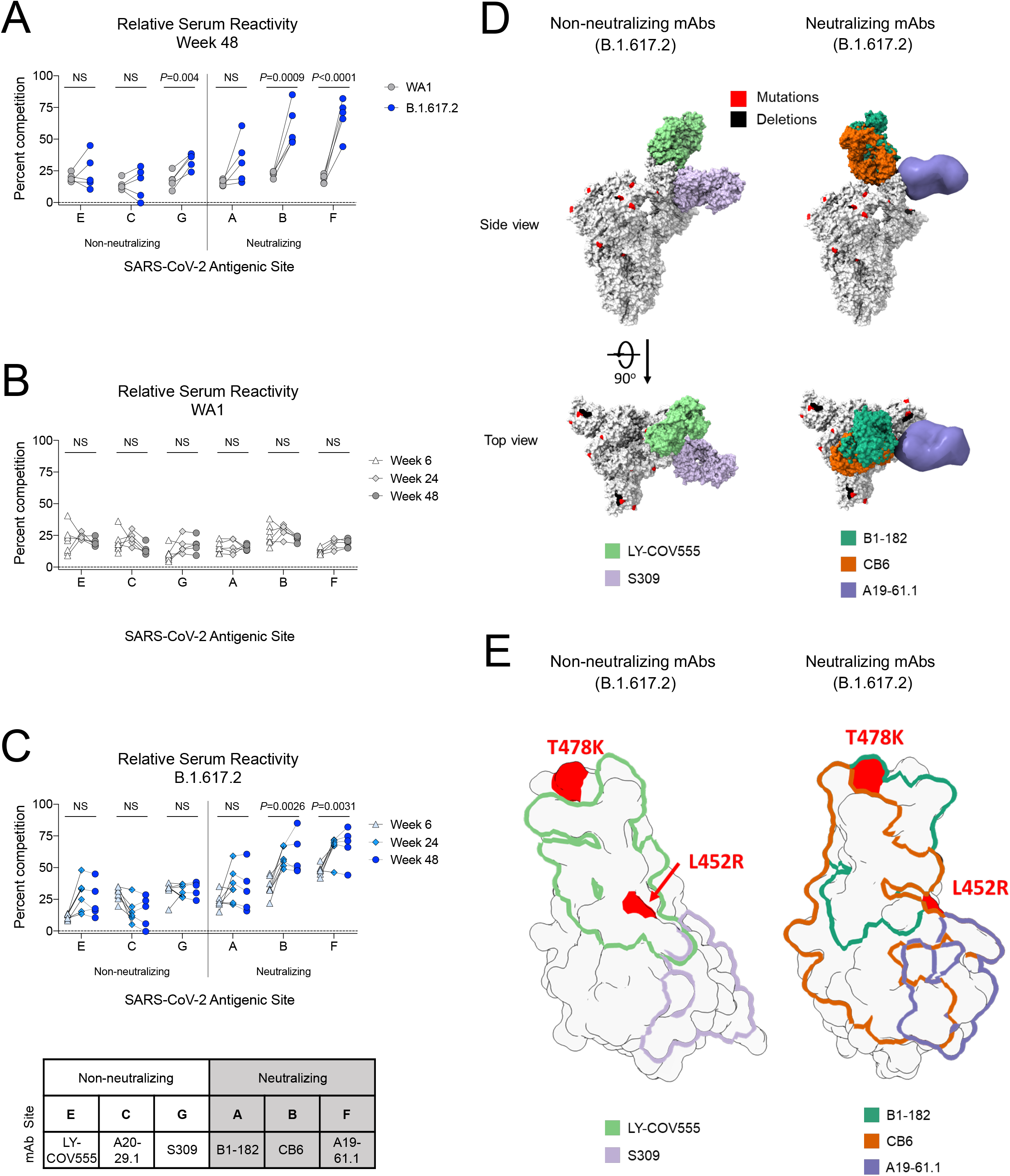
B.1.617.2 S-2P-binding serum antibodies recognize epitopes associated with neutralization. (A) Relative serum reactivity was measured as percent competition of total measured serum antibody S-2P binding competed by single monoclonal antibody (mAb) targeting cross-reactive RBD epitopes on both WA1 and B.1.617.2 S-2P at Week 48 post-immunization. Antigenic sites are defined by mAbs LY-COV555 (site E), A20-29.1 (site C), S309 (site G), B1-182 (site A), CB6 (site B), and A19-61.1 (site F). 5 NHP per group. Statistical analysis shown for percent competition of binding to indicated epitopes on WA1 S-2P in comparison to B.1.617.2 S-2P. (B-C) Longitudinal analysis of relative serum reactivity to cross-reactive RBD epitopes on both WA1 (B) and B.1.617.2 S-2P (C) was evaluated at 6, 24 and 48 weeks post-immunization. 5-8 NHP per group. Statistical analysis shown for percent competition of binding to indicated epitopes at week 48 in comparison to week 6. (D) SARS-CoV-2 S models with B.1.617.2 mutations indicated in red and deletions in black shown in complex with non-neutralizing (LY-COV555, S309) and neutralizing (B1-182, CB6, A19-61.1) mAbs. (E) Footprints of both non-neutralizing (LY-COV555, S309) and neutralizing (B1-182, CB6, A19-61.1) mAbs indicate areas of binding on B.1.617.2 receptor binding domain (RBD) with mutations highlighted in red. See also Table S2 for Barnes classifications of mAbs.

In contrast, longitudinal analysis of serum reactivity to cross-reactive epitopes on B.1.617.2 S-2P showed significantly increased relative reactivity to antigenic sites B and F (represented by neutralizing mAbs CB6 and A19-61.1) from week 6 to 48 (B: *P*=0.0026; F: *P*=0.0031) (Fig 2C). Structural models show different angles of approach between non-neutralizing (LY-COV555, S309) and neutralizing (B1-182, CB6, A19-61.1) mAbs in complex with B.1.617.2 S-2P (Fig 2D). Modeling of binding footprints on B.1.617.2 RBD show immuno-focusing occurs on neutralizing epitopes with areas of binding outside mutations T478K and L452R (CB6 and A19-61.1) (Fig 2E). Additionally, increased reactivity to B.1.617.2 S-2P as compared to WA1 S-2P (Fig 2A) was only observed to epitopes binding outside RBD mutations (T478K, L452R) (Fig 2E). These data indicate that although serum antibodies targeting B.1.617.2 S-2P decreased over time (Fig 1C), the proportion of serum antibodies targeting epitopes associated with neutralization increased, suggesting a maturing of the humoral immune response and immuno-focusing on conserved epitopes associated with neutralization against B.1.617.2.

### Kinetics of B.1.617.2 RBD-binding IgG and IgA antibodies in the upper and lower airway

Antibodies are likely to provide the initial immune mechanism to control viral replication in the upper and lower airways (Corbett et al., 2021b; Francica et al., 2021; Froberg et al., 2021; Gilbert et al., 2021; Khoury et al., 2021; Roozendaal et al., 2021). To determine the persistence of antibody titers in these relevant anatomical sites, we collected bronchoalveolar lavage (BAL) fluid and nasal washes at weeks 6, 25 and 42. WA1 RBD IgG binding titers in the BAL were highest at week 6 and declined by week 42 such that GMT were almost 5000-fold less than at week 6 (*P*=0.0007) (Fig. 3A). B.1.617.2 RBD-binding IgG titers were lower in the BAL, with a 6-fold GMT reduction compared to WA1 at week 6 and a 4-fold reduction at week 42 (Fig. 3C and Fig. S2A). B.1.351 RBD-binding titers were lowest, with a 342-fold and 57-fold reduction compared to WA1 at weeks 6 and 42, respectively (Fig. S2C). In addition, B.1.617.2 and B.1.351 RBD-binding titers followed a similar trend as WA1 RBD-binding titers with all antibody levels decreasing by week 42 (B.1.617.2: *P*=0.0008; B.1.351: *P*=0.0007). In contrast to the decreased antibody responses in BAL over time, WA1, B.1.351 and B.1.617.2 RBD-binding titers in the nasal washes were highest at week 25 and remained stable by week 42 (Week 6 vs Week 42: *P*>0.05) (Fig. 3B, D and Fig. S2B, D).

**Figure 3.**
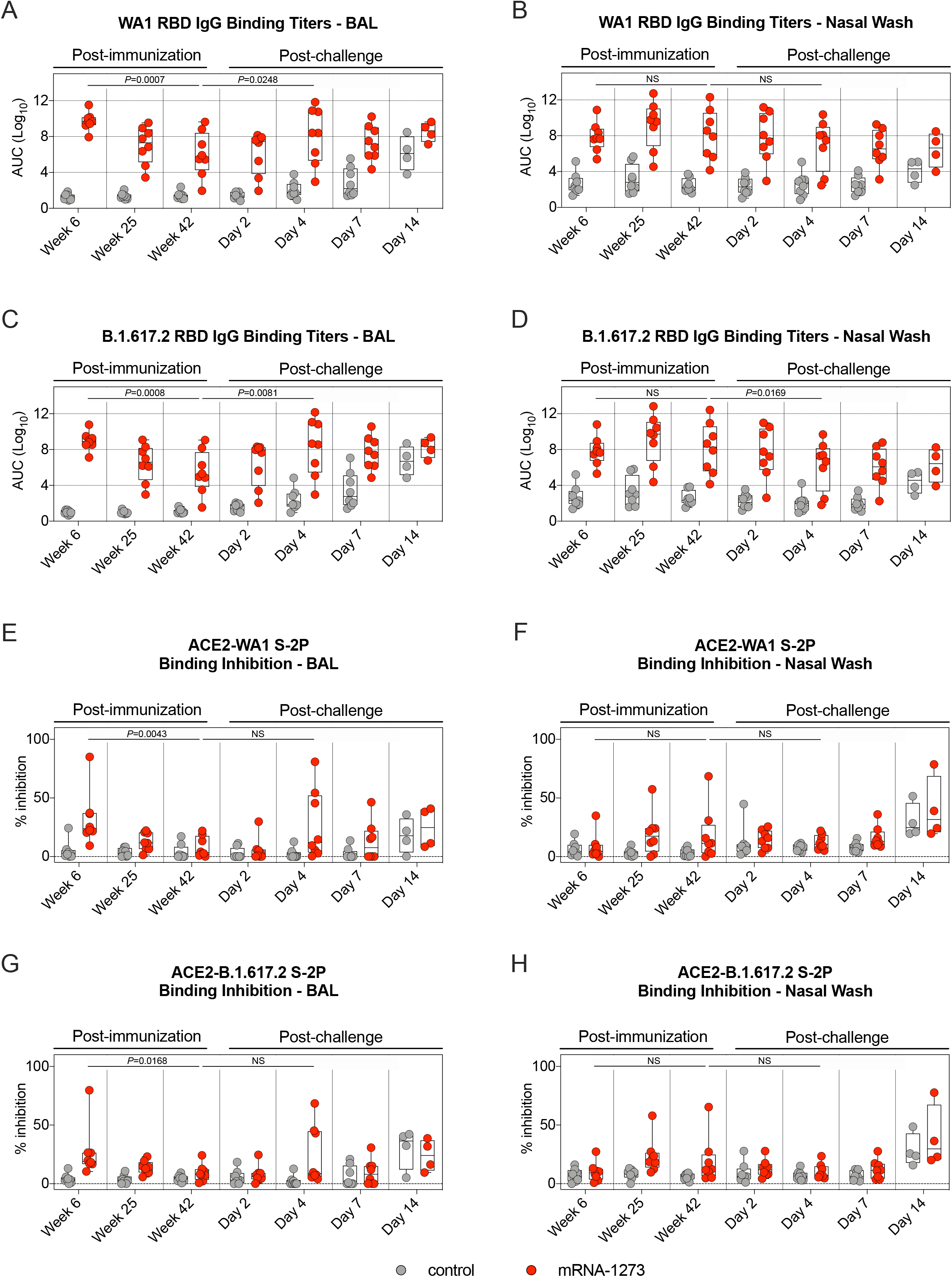
RBD-binding mucosal antibodies observe distinct kinetic patterns in the upper and lower airway. (A-D) BAL and nasal washes were collected at weeks 6, 25 and 42 post-immunization, and days 2, 4, 7 and 14 post-challenge. (A-B) WA1 and (C-D) B.1.617.2 RBD-binding IgG titers in the lower (A, C) or upper airway (B, D). Circles in (A-D) indicate individual NHP. Boxes represent interquartile range with the median denoted by a horizontal line. Dotted lines are for visualization purposes and denote 4-log_10_ increases in binding titers. 4-8 NHP per group. (E-H) BAL and nasal washes were collected at weeks 6, 25 and 42 post-immunization, and days 2, 4, 7 and 14 post-challenge. All samples diluted 1:5. SARS-CoV-2 WA1 (E-F) and B.1.617.2 (G-H) S-2P binding to ACE2 measured both alone and in the presence of BAL (E, G) or nasal wash (F, H) in order to calculate % inhibition. Circles denote individual NHP. Boxes represent interquartile range with the median denoted by a horizontal line. Dotted lines set to 0% inhibition. 4-8 NHP per group. Statistical analysis in (A-H) shown for mRNA-1273 cohort only. See also Figure S2 for IgA responses and kinetics of B.1.617.2 RBD-binding IgG titers for individual NHP.

SARS-CoV-2 RBD-binding IgA titers in nasal washes and BAL were also assessed (Fig. S2E-H). At week 6, GMT in the BAL to WA1 RBD and B.1.617.2 RBD were 9-fold and 26-fold higher in vaccinated NHP than in controls, respectively. IgA titers were indistinguishable from controls by week 24 in the BAL. IgA titers in the nasal washes of vaccinated NHP were similar to those from controls at all timepoints.

In our previous NHP studies, we have been unable to detect neutralizing antibodies in the upper or lower airway following mRNA-1273 immunization using either pseudovirus or live virus assays. Here we used the S-2P-ACE2 binding inhibition assay which provides a highly sensitive assessment of antibody function to extend the analysis of binding titers in the airways. Consistent with the findings from the sera (Fig. S1B-C), the highest S-2P-ACE2 binding in BAL was detected at week 6 and declined by week 42 (WA1: *P*=0.0043; B.1.617.2: *P*=0.0168) (Fig. 3E, G). By contrast, S-2P-ACE2 binding inhibition in the upper airway was not statistically different between weeks 6 and 42 (Fig. 3F, H). Together, these results suggest that the kinetics and durability of antibody responses in the upper respiratory tract (nasal washes) are distinct from those in the blood or lungs (BAL).

### Kinetics of S-specific memory B cells responses

Sustained antibody production and increased secondary responses following a vaccine boost or viral challenge are dependent on antigen-specific memory B cell responses (Gaebler et al., 2021; Goel et al., 2021a). A number of human studies have shown that mRNA S-2P vaccines and/or SARS-CoV-2 infection induce S-specific memory B cells that persist over time (Dan et al., 2021; Goel et al., 2021b; Rodda et al., 2021; Turner et al., 2021; Wang et al., 2021d; Wang et al., 2021e). Here, the frequency of WA1 S-and B.1.617.2 S-specific memory B cells following mRNA-1273 vaccination in NHP were assessed by flow cytometry using fluorescent probes (Fig. S3A). At week 6, a median value of 2.5% of all memory B cells were dual-specific for WA-1 and B.1.617.2 in comparison to 0.14% for WA-1 and 0.09% for B.1.617.2 S-specific alone (Fig. 4A-D). Although the total percentage of probe-binding memory B cells decreased through week 42, the high frequency of dual-binding to single-binding cells remained constant, with a geometric mean proportion of dual-binding cells greater than 85% at all timepoints (Fig. 4E). These data are consistent with recent reports that mRNA-1273 and BNT162b2 elicit S-specific memory B cells that are capable of binding both the vaccine-matched strain and VOC (Corbett et al., 2021a; Goel et al., 2021b). The memory phenotype of these cells was also determined over time after vaccination (Fig. S3B). Over the course of one year, S-2P-binding B cells shifted from 87% having an activated memory phenotype to a more balanced distribution of 43% activated memory, 32% resting memory, and 15% tissue-like memory cells (Fig. 4F). Together, these data suggest that mRNA-1273 induces durable, broad cross-strain B cell memory to SARS-CoV-2.

**Figure 4.**
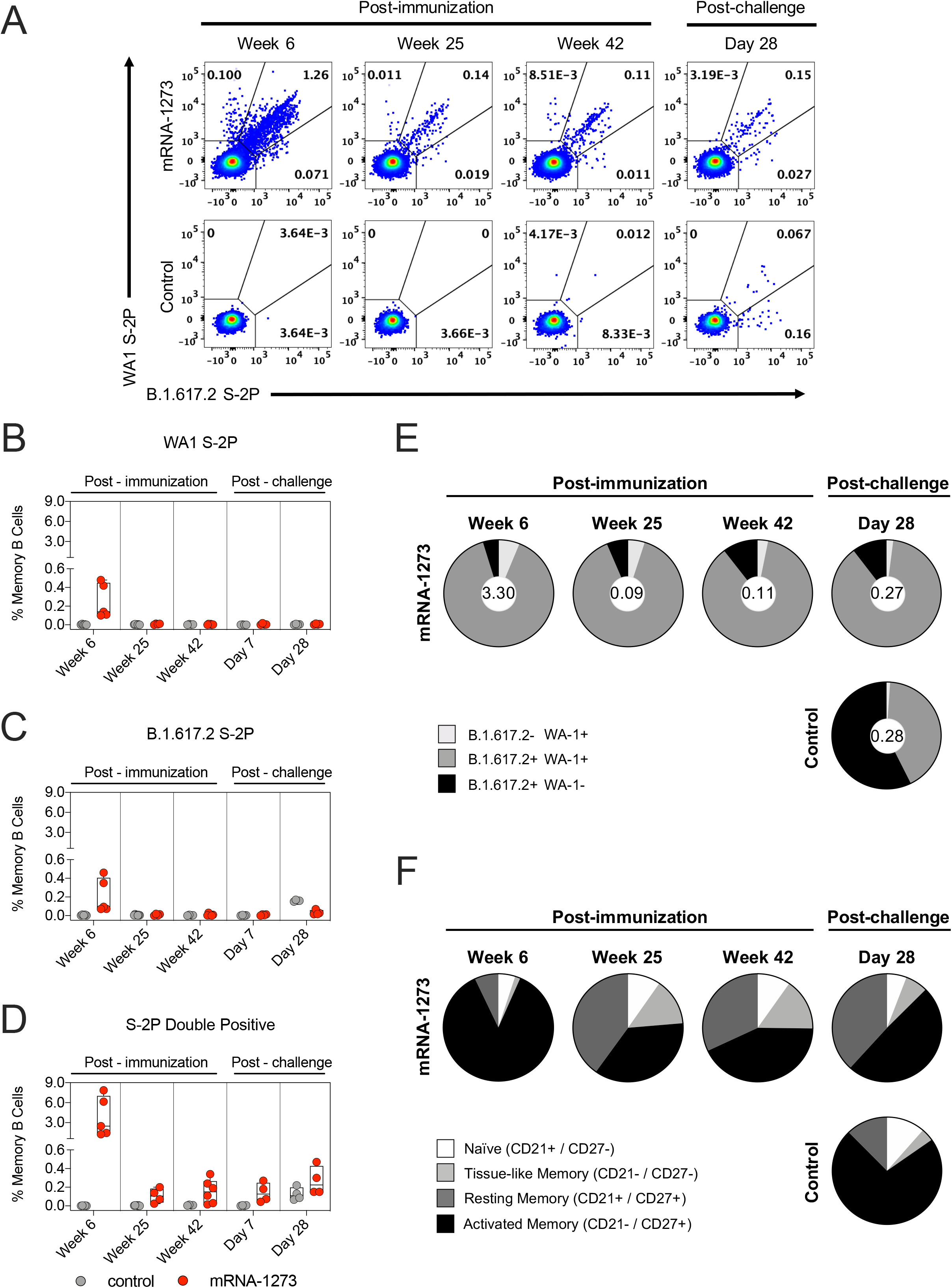
mRNA-1273 elicits memory B cells that bind both WA1 and B.1.617.2. (A) Representative flow cytometry plots showing WA1 S-2P-specific, B.1.617.2 S-2P-specific, and dual-binding memory B cells at weeks 6, 25 and 42 post-immunization, and day 28 post-challenge. *Top row.* mRNA-1273 group. *Bottom row.* mRNA control group. (B-D) Percentage of all memory B cells that are specific for WA1 S-2P (B), B.1.617.2 S-2P (C), or both (D) at weeks 6, 25 and 42 post-immunization, and days 7 and 28 post-challenge. Circles in (B-D) indicate individual NHP. Boxes represent interquartile range with the median denoted by a horizontal line. 4-7 NHP per group. Break in Y-axis indicates a change in scale without a break in the range depicted. (E) Pie charts indicating the proportion of SARS-CoV-2 S-specific memory B cells that bind to WA1 S-2P alone (light gray), B.1.617.2 S-2P alone (black), or both (dark gray) at weeks 6, 25 and 42 post-immunization, and day 28 post-challenge. Values inside pie charts represent the percentage of all memory B cells that bind any SARS-CoV-2 S-2P probe at each timepoint. 4-7 NHP per group. *Top row.* mRNA-1273 group. *Bottom row.* mRNA control group. (F) Pie charts indicating the proportion of SARS-CoV-2 S-specific B cells with a phenotype consistent with naïve (white), tissue-like memory (light gray), resting memory (dark gray), or activated memory (black) cells at weeks 6, 25 and 42 post-immunization, and day 28 post-challenge. 4-7 NHP per group. *Top row.* mRNA-1273 group. *Bottom row.* mRNA control group. See also Figure S3 for gating strategy.

### mRNA-1273 induces T_H_1 and T_FH_ responses

Our previous reports showed that mRNA-1273 immunization elicits SARS-CoV-2 S-specific T_H_1, CD40L^+^ T_FH_, and IL-21^+^ T_FH_ cells which decrease over a 6-month period in NHP (Corbett et al., 2021a). Here, S-specific T cell responses were assessed through week 42 in blood and BAL (Fig. S3C). Consistent with our prior studies, T_H_1 but not T_H_2 responses were detected in the blood at week 6, decreasing by week 25. T_H_1 levels were low to undetectable by week 42 (Week 6 vs Week 42: *P*=0.0261) (Fig. 5A-B). mRNA-1273 induced both CD40L^+^ and IL-21^+^ T_FH_ cells at week 6. By week 42, both populations had declined (CD40L^+^: *P*=0.0164; IL-21^+^: *P*=0.0166) (Fig. 5D-E). Median CD8^+^ T cell responses in the blood were low (Fig. 5C), although we did detect a low frequency of CD8^+^ T cells in the BAL at week 6 (Fig. 5H).

**Figure 5.**
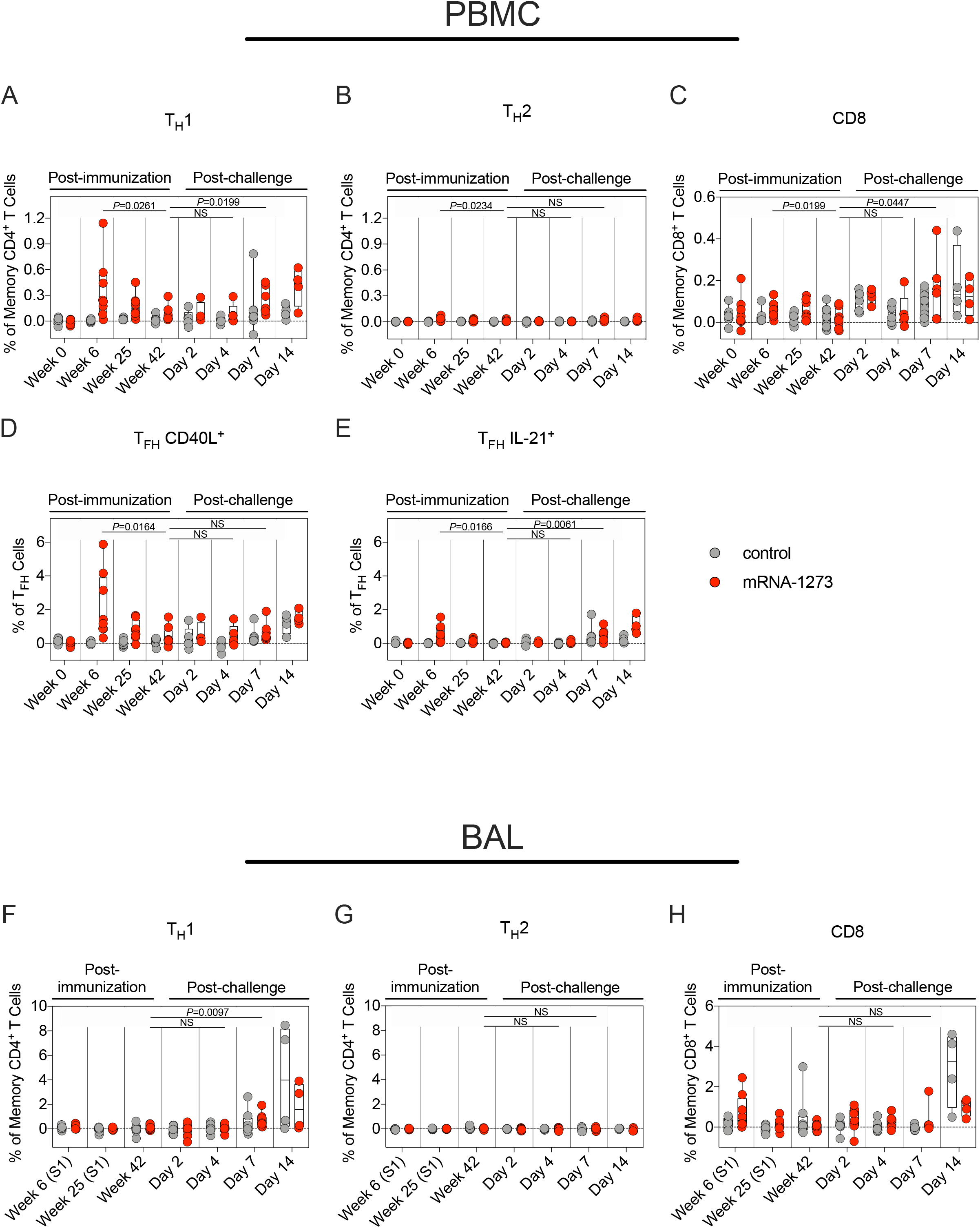
B.1.617.2 challenge induces an anamnestic T cell response in the periphery of vaccinated NHP. (A-E) PBMC collected at weeks 0, 6, 25 and 42 post-immunization, and days 2, 4, 7 and 14 post-challenge. Cells were stimulated with SARS-CoV-2 S1 and S2 peptide pools and then measured by intracellular staining. (A-B) Percentage of memory CD4^+^ T cells with (A) T_H_1 markers (IL-2, TNF or IFNγ) or (B) T_H_2 markers (IL-4 or IL-13) following stimulation. (C) Percentage of CD8^+^ T cells expressing IL-2, TNF or IFNγ. (D-E) Percentage of T_FH_ cells that express (D) CD40L or (E) IL-21. (F-H) BAL fluid was collected at weeks 6, 25 and 42 post-immunization, and days 2, 4, 7 and 14 post-challenge. Lymphocytes in the BAL were stimulated with S1 and S2 peptide pools and responses measured by intracellular cytokine staining using T_H_1 (F), T_H_2 (G), and CD8 markers (H). Samples from weeks 6 and 25 were only stimulated with S1 peptides. Circles in (A-H) indicate individual NHP. Boxes represent interquartile range with the median denoted by a horizontal line. Dotted lines set at 0%. Reported percentages may be negative due to background subtraction. 4-8 NHP per group. Statistical analysis shown for mRNA-1273 cohort only. See also Figure S3 for gating strategy and Figure S4 for T cell responses to N peptides in the lower airway.

### Protection Against B.1.617.2 one year after mRNA-1273 Vaccination

For the viral challenge, an isolate of B.1.617.2 was obtained after one passage and the canonical S mutations associated with B.1.617.2, including T19R, L452R, T478K, D614G, P681R, as well as the NTD deletion (Fig. S5A) were verified by sequencing. This viral stock was first tested in hamsters (Fig. S5C) and nonhuman primates (Fig. S5D-E) to confirm its pathogenicity and define a dose for challenge.

mRNA-1273 and mRNA control NHP were challenged 49 weeks after the initial immunization with 2×10^5^ plaque forming units (PFU) of virus via intratracheal (IT) and intranasal (IN) routes (Fig. S1A). BAL and nasal swabs (NS) were collected on days 2, 4, 7 and 14 following challenge and qRT-PCR was used to measure viral replication by assessing sgRNA copies encoding for the SARS-CoV-2 E and N genes. On day 2, geometric mean sgRNA_E copies in the lower airway of controls and vaccinated NHP were 1×10^6^ and 9×10^4^ per mL, respectively (*P*>0.05) (Fig. 6A). sgRNA_E copies in the lower airway of vaccinated NHP declined rapidly over the following days, with geometric mean copies of 9×10^2^ on day 4 and 1×10^2^ on day 7. sgRNA_E copies in the lungs of unvaccinated NHP remained significantly elevated compared to vaccinated NHP, with a geometric mean of 1×10^5^ on day 7 (mRNA-1273 vs control: *P*<0.0001). These results differed from our earlier findings that NHP vaccinated with 100µg mRNA-1273 and then challenged only 4-8 weeks later largely controlled WA1 or B.1.351 replication in the lower airway by day 2 (Corbett et al., 2020; Corbett et al., 2021c).

**Figure 6.**
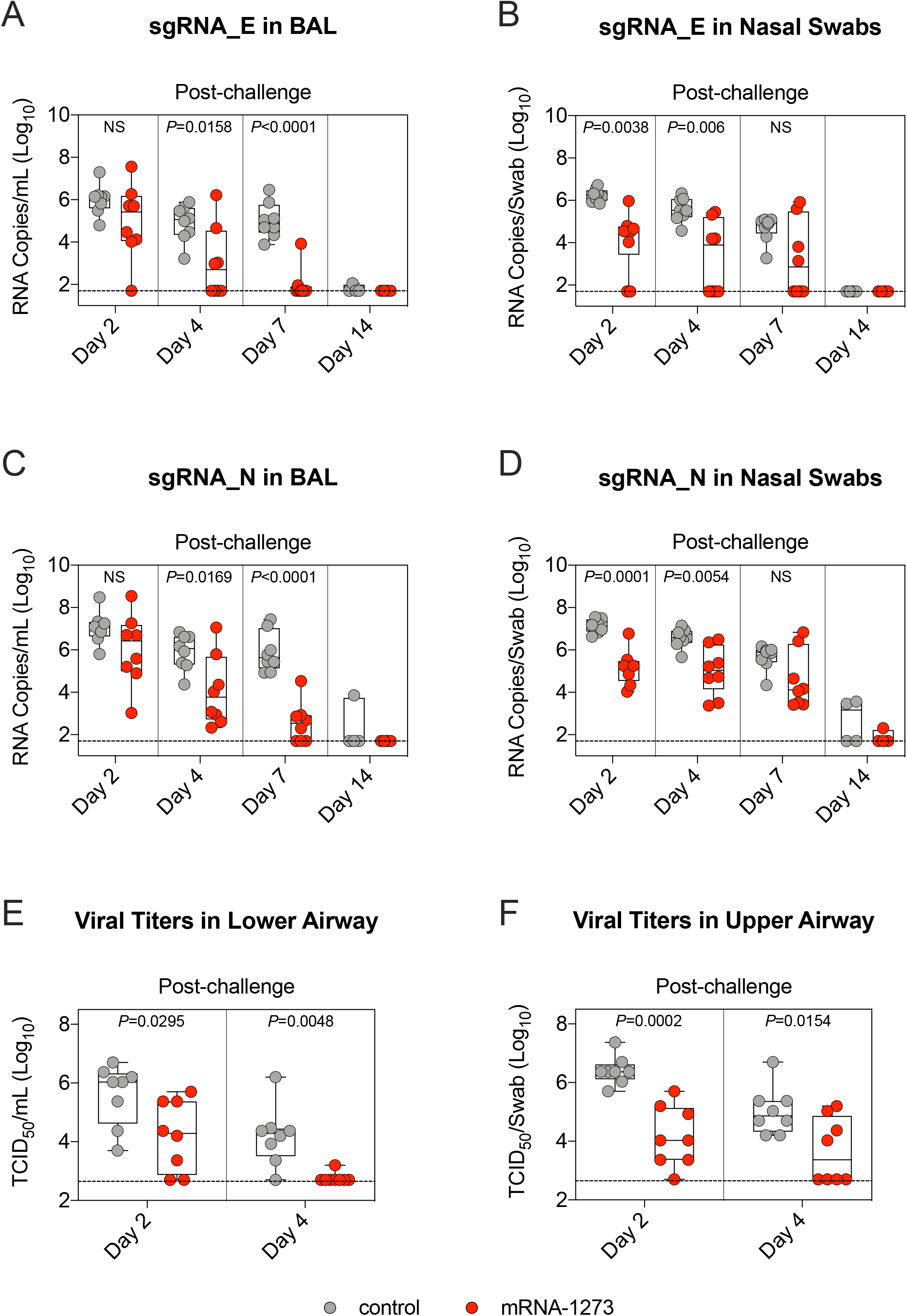
B.1.617.2 replication is reduced in the upper and lower airway of mRNA-1273-immunized NHP. (A-D) BAL and NS were collected 2, 4, 7 and 14 days after challenge with 2×10^5^ PFU B.1.617.2. Copy numbers of sgRNA_E (A-B) and sgRNA_N (C-D) in the BAL (A, C) or nose (B, D). 4-8 NHP per group. (E-F) Viral titers per mL of BAL (E) or per NS (F) collected 2 and 4 days after challenge. 8 NHP per group. Circles in (A-F) indicate individual NHP. Boxes represent interquartile range with the median denoted by a horizontal line. Dotted lines indicate assay limit of detection. Statistical analysis shown for mRNA-1273 cohort in comparison to control NHP at indicated timepoints. See also Figure S5 for B.1.617.2 challenge stock characterization, Figure S6 and Table S3 for correlates analysis, and Figure S7 for non-specific immune responses following challenge.

In contrast to the lower airway, we observed less reduction in viral replication in the upper airway following mRNA-1273 immunization (Fig. 6B). On day 2, geometric mean copies of sgRNA_E were 2×10^6^ and 6×10^4^ per NS of controls and vaccinated NHP, respectively (*P*=0.0038). On days 4 and 7, copies declined to 2×10^3^ and 1×10^3^ for the mRNA-1273 group and 4×10^5^ and 5×10^4^ in the control group (Day 4: *P*=0.006; Day 7: *P*>0.05). By day 7, 6/8 and 4/8 vaccinated NHP had undetectable sgRNA_E levels in the lower airway and upper airway, respectively. All control NHP still had detectable sgRNA_E copies at that time point. We observed similar trends with sgRNA_N (Fig. 6C-D), but with higher geometric mean copy numbers in agreement with our prior publications (Corbett et al., 2021b; Corbett et al., 2021c).

We euthanized 4/8 NHP in each cohort on day 7 for assessment of lung pathology but continued to monitor the remaining animals for an additional week. On day 14, 1/4 and 2/4 control NHP had detectable sgRNA_N in the lower and upper airway, respectively. In contrast, 0/4 and 1/4 vaccinated NHP had detectable sgRNA_N in the same compartments. It is noteworthy that the only animal in the mRNA-1273 cohort with detectable sgRNA_N at day 14 had undetectable pseudovirus neutralizing antibody titers to both D614G and B.1.617.2 at week 48 (Fig. 1F).

To provide an additional assessment of protection, we measured tissue culture infectious dose (TCID_50_) in samples taken from the lungs and nose on days 2 and 4 (Fig. 6E-F). By day 4, only 1/8 animals in the mRNA-1273 group had detectable virus in the lower airway compared to 7/8 in the control mRNA group (*P*=0.0048). In the upper airway, 4/8 vaccinated animals had detectable virus, while all control animals had detectable virus (*P*=0.0154).

We previously established binding and neutralizing antibody titers in the sera as an immune correlate of protection from WA1 and B.1.351 replication in NHP, when both challenges were given 4-8 weeks after boost (Corbett et al., 2021b; Corbett et al., 2021c). Here, when B.1.617.2 challenge was approximately one year after mRNA-1273 vaccination, we looked to see whether these associations held between viral replication and serum WA1 S-2P binding titers (WHO units), B.1.617.2 RBD-binding IgG titers, D614G lentiviral pseudovirus neutralizing titers, and B.1.617.2 lentiviral pseudovirus neutralizing titers at peak (week 6) and immediately prior to challenge (week 48) (Fig. S6 and Table S3). Unlike in the challenges that were conducted soon after vaccination, none of these measurements correlated with viral replication in the lower airway. However, neutralizing titers at week 48 were significantly inversely correlated with viral replication in the upper airway at day 2 after challenge (D614G: *P*=0.0149; B.1.617.2: *P*=0.0446) but not on any subsequent days (Fig. S6G-H).

### B.1.617.2 challenge elicits an anamnestic antibody response in the lower airway

Analysis of immune responses after challenge provides data on the kinetics of anamnestic responses which may reveal potential mechanisms of protection. First, IgG binding titers and neutralizing antibody titers were assessed in sera at days 2, 4, 7 and 14 following challenge. We observed a clear primary response by day 14 in control NHP, with increased binding to WA1, B.1.351, and B.1.617.2 RBDs (Fig. 1B-D). B.1.617.2 GMT increased 37000-fold relative to week 48 and was higher than WA1 GMT at day 14. In contrast, B.1.617.2 GMT in the mRNA-1273 group increased only 490-fold at day 14 relative to week 48 and was equivalent to WA1 GMT at that time. In addition, WA1 S-2P binding GMT in control NHP increased 39-fold from 0.9 BAU/mL to 35 BAU/mL between week 48 after immunization and day 14 after challenge. The corresponding increase in vaccinated NHP was smaller, rising 5-fold from 189 to 1030 BAU/mL (Fig. 1A).

Pseudovirus neutralizing responses were also measured to D614G and B.1.617.2 at days 2, 4, 7 and 14. The hierarchy of neutralizing antibody titers elicited by the vaccine remained stable until day 14, when titers to B.1.617.2 approached those of D614G (*P*>0.05) (Fig. 1F). These findings show that while serum antibody responses did not increase at a faster rate in vaccinated NHP than in controls, they were boosted by infection.

We next measured mucosal antibody levels in the upper and lower airway. RBD-binding IgG titers in the upper airways of vaccinated animals showed a modest decline (Fig. 3D and Fig. S2B) despite virus persistence in the nose (Fig. 6B, D, F), which may reflect clearance of viral protein-bound antibodies. We detected a strikingly different trend in the lower airway, however. B.1.617.2-binding GMT in the control group increased slowly relative to week 42 over the post-challenge observational period, rising 18-fold and 213-fold on days 4 and 7 respectively. The antibody binding response to B.1.617.2 in vaccinated NHP was more rapid, rising 590-fold on day 4 before falling to 157-fold on day 7 compared to week 42 (Fig. 3C). Antibody titers to all 3 variants were significantly greater 4 days after challenge than 42 weeks after immunization in the mRNA-1273 vaccinated group (WA1: *P*=0.0248; B.1.617.2: *P*=0.0081; B.1.351: *P*=0.0038) (Fig. 3A, C and Fig S2). Last, we did not detect differential IgA responses or ACE2 binding inhibition between vaccinated animals or controls in either the lower or upper airway (Fig. 3E-H and Fig S2). These data show that the mRNA-1273-vaccinated NHP made an anamnestic IgG response in the lower airways.

Although the total IgG concentration in the lungs of both vaccinated and unvaccinated NHP increased following challenge, this increase was greater in the control cohort. On days 2, 4, 7 and 14 after challenge, geometric mean IgG concentrations were 1.5-to 3-fold lower in the vaccinated cohort than in the control group (Fig. S7A). This rapid response after challenge is consistent with our prior experience of a protein SARS-CoV-2 vaccine in which we observed a rapid increase in both total IgG and measles morbillivirus (MeV) titers in primates that had also been vaccinated against MeV (Francica et al., 2021). In the present study, although binding titers were similar between both cohorts prior to challenge, responses to MeV increased more rapidly in the control group. On day 4, GMT to MeV in the lower airway were 3.3-fold greater in the control animals than in the mRNA-1273 group (Fig. S7B).

An anamnestic S-specific B cell response in the lower airways would likely be the underlying mechanism for an increase in local antibody titers. While we were unable to collect a sufficient number of B cells in BAL to analyze after B.1.617.2 challenge, we were nevertheless able to interrogate these cells in blood. Memory B cells which bound both WA1 S-2P and B.1.617.2 S-2P dominated the S-specific immune response of vaccinated NHP. While the total frequency of all S-binding memory B cells increased, the proportion of these dual-specific cells remained between 85-90% (Fig. 4E). In contrast, B.1.617.2 challenge in control NHP elicited a higher frequency of memory B cells specific only for B.1.617.2 S, with a geometric mean frequency of 57% at day 28 (Fig 4E). 72% of S-binding B cells in control NHP had a phenotype consistent with activated memory B cells, in contrast to the vaccinated cohort in which 49% of cells had an activated memory phenotype and 38% of cells had a resting memory phenotype (Fig. 4F).

### T cell responses in blood and BAL following vaccination and B.1.617.2 challenge

After showing a potential role for memory B cell responses in controlling virus replication in lower airways, we also measured T cell responses to SARS-CoV-2 S in the blood and BAL following challenge. By day 7, T_H_1 responses were elevated as compared to week 42 in both compartments (Serum: *P*=0.0199; BAL: *P*=0.0097). However, the range of S-specific T_H_1 CD4^+^ T cell frequencies in the BAL of control and vaccinated NHP overlapped (Fig. 5A, F). While we observed increased IL-21^+^ T_FH_ responses by day 7 (*P*=0.0061) (Fig. 5E), we did not detect any T_H_2 responses following challenge in vaccinated or control animals (*P*>0.05) (Fig. 5B, G). In contrast to the minimal vaccine-elicited CD8^+^ T cells prior to challenge, both control and vaccinated NHP mounted CD8^+^ T cell responses following challenge (Fig. 5C, H). Finally, T_H_1 and CD8^+^ responses in the lungs to SARS-CoV-2 N, which was not encoded by mRNA-1273, were only present in control NHP (Fig. S4). Together, these data demonstrate that vaccination with mRNA-1273 not only elicited B and T cellular responses to SARS-CoV-2 S following challenge but also protected NHP from encountering sufficient levels of SARS-CoV-2 N to mount a primary response to that antigen.

### mRNA-1273 protects the lower airway from severe inflammation

Because sufficient virus was present in the lower airway of vaccinated NHP to elicit both a local anamnestic antibody response and increased S-specific B and T cell populations, we assessed if mRNA-1273 vaccination protected the lungs from gross pathologic changes. Lung samples from 4/8 NHP in each group were evaluated for pathology and detection of viral antigen (VAg) 7 days after B.1.617.2 challenge. SARS-CoV-2 VAg (red arrowheads) was detected in the lungs of 4/4 control animals (Fig. 7B, C) and was not detected in any of the animals in the vaccinated cohort (Fig. 7A, C). Inflammation ranged from minimal to moderate across lung samples from animals that received mRNA-1273 and from minimal to severe in control NHP (Fig. 7C). The inflammatory changes in the lungs of vaccinated NHP were characterized by a mixture of macrophages and polymorphonuclear cells present within some alveolar spaces and mild to moderate expansion of alveolar capillaries with mild Type II pneumocyte hyperplasia (Fig. 7A). Changes in control NHP were more consistent with lymphocytes, histiocytes and fewer polymorphonuclear cells associated with more prominent and expanded alveolar capillaries, collections of cells with alveolar air spaces, occasional areas of perivascular and peribronchiolar inflammation, and Type II pneumocyte hyperplasia (Fig. 7B). The protection observed in the lungs suggest that although binding and neutralizing antibody titers had declined one year after vaccination (Fig. 1), leaving the lower airway susceptible to virus replication in the first two days after challenge (Fig. 6), a local anamnestic response to antigen previously encountered in the vaccine proved sufficient to prevent severe disease.

**Figure 7.**
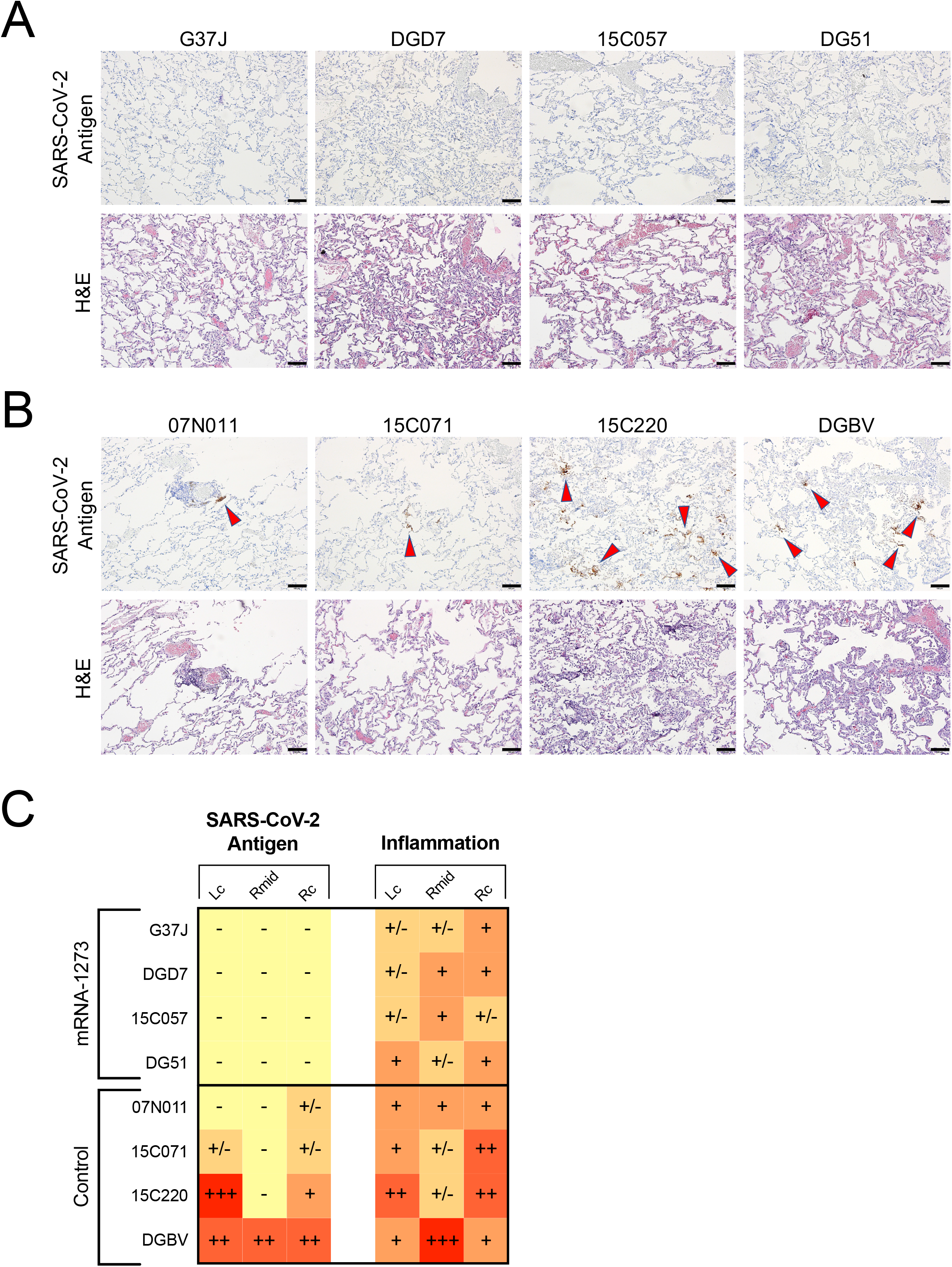
mRNA-1273 provides durable protection in the lower airway from B.1.617.2. (A-B) Representative images of lung samples 7 days after B.1.617.2 challenge from 4 NHP that received mRNA-1273 (A) or mRNA control (B). *Top row.* Detection of SARS-CoV-2 antigen by immunohistochemistry with a polyclonal anti-N antibody. Antigen-positive foci are marked by red arrows. *Bottom row.* Hematoxylin and eosin staining (H&E) illustrating the extent of inflammation and cellular infiltrates. Images at 10x magnification with black bars for scale (100µm). (C) SARS-CoV-2 antigen and inflammation scores in the left cranial lobe (Lc), right middle lobe (Rmid), and right caudal lobe (Rc) of the lungs 7 days after B.1.617.2 challenge. Antigen scoring legend: **-** no antigen detected; **+/-** rare to occasional foci; **+** occasional to multiple foci; **++** multiple to numerous foci; **+++** numerous foci. Inflammation scoring legend: - minimal to absent inflammation; **+/-** minimal to mild inflammation; **+** mild to moderate inflammation; **++** moderate to severe inflammation; **+++** severe inflammation. Horizontal rows correspond to individual NHP depicted above (A-B).

## DISCUSSION

mRNA-1273 vaccine efficacy against both symptomatic and asymptomatic infection with B.1.617.2 is reduced compared to ancestral strains, a result of both variant-specific mutations which confer diminished sensitivity to neutralization and waning antibody titers over time following vaccination (Baden et al., 2021b; Bruxvoort et al., 2021; Choi et al., 2021; Puranik et al., 2021; Tada et al., 2021). Here, we studied protection against B.1.617.2 infection by mRNA-1273 almost one year after vaccination in NHP. Our principal findings were: (1) protection in the lower airway was complete by day 4 but delayed compared to prior NHP studies in which viral challenges were done much closer to the time of vaccination; (2) anamnestic antibody responses were detected in BAL at day 4 in vaccinated animals; (3) and binding and neutralizing antibody titers in blood and BAL decreased over 48 weeks.

Prior studies by us and others in NHP (Corbett et al., 2021a) and humans (Doria-Rose et al., 2021; Goel et al., 2021b) show reduction of neutralizing antibodies in sera over 6 months following mRNA vaccination. Here, antibody binding and neutralizing titers also decline in the sera over one year, and we extend these data by showing similar findings in BAL suggesting that antibody measurements in serum may be a surrogate for such responses in the lung. By contrast, antibodies remain relatively stable from week 24-48 in the upper airway. The IgG stability in the nasal tract is notable considering the source of this immunoglobulin class is primarily transudation from the blood (Renegar et al., 2004). These data are consistent with a prior study showing that subcutaneous vaccines elicit antibodies that are detectable in the nose for at least 6 months (Clements and Murphy, 1986). The differential kinetics for the stability of IgG responses in the upper respiratory and lower respiratory tract suggest unique mechanisms of clearance within these different anatomical sites.

The limited reduction in viral replication in the lower airway 2 days after challenge in this present study stands in contrast to prior studies in which mRNA-1273-vaccinated NHP were challenged 4-8 weeks after vaccination and lower airway viral replication was largely controlled by day 2 (Corbett et al., 2020; Corbett et al., 2021c). The decline of antibody titers in the lungs over one year, which are replete with epithelial target cells, may explain the different kinetics in virus control. Importantly, vaccinated NHP had a striking anamnestic antibody response in the lower airways, with a 590-fold increase in B.1.617.2 GMT by day 4 after challenge compared to the pre-challenge timepoint, and there was no detectable sgRNA_E in 6/8 vaccinated NHP by day 7. This suggests that protection in the lower airway is durable but somewhat delayed and may be dependent on a recall antibody response. It is also notable that while we were able to detect binding and some functional antibodies in the blood and mucosa through the S-2P-ACE2 inhibition assay at time of challenge, neutralizing titers to B.1.617.2 were low to undetectable. Thus, vaccine-elicited protection in the lower airway could be mediated by neutralizing antibodies below our limit of detection or through Fc-mediated functions, which has been shown to be important for protection by mAb treatment (Winkler et al., 2021). It is also noteworthy that although the quantity of antibodies had decreased one year after vaccination, the quality was improved as evidenced by the increase in avidity and the shift in epitope dominance of B.1.617.2 S-binding antibodies towards sites associated with neutralization. In the nose, the 1-log_10_ reduction in sgRNA_N copy numbers on days 4 and 7 was consistent with what we have observed after our prior B.1.351 challenge with the same dose and regimen of mRNA-1273 (Corbett et al., 2021c) and shows that there is a higher threshold of antibody required for protection in the upper airway compared to the lower airway (Corbett et al., 2021b). These data are also consistent with studies in human vaccinees showing that there can be a significantly higher level of protection against severe disease than mild or asymptomatic infection (Baden et al., 2021b; Bruxvoort et al., 2021; Puranik et al., 2021; Tang et al., 2021).

Rapid control of virus replication in both upper and lower airways has potential implications for transmission. Smaller aerosols (<20 microns) enriched with virus are generated mainly in the lower airways and their inhalation may lead to transmission (Wang et al., 2021a). Therefore, in the context of limited virus control in the upper airway and slower kinetics of control in the lower airway, a boost to increase antibody titers may eventually be warranted.

Antibodies elicited by mRNA-1273 are capable of binding multiple VOC in NHP and humans (Corbett et al., 2021a; Goel et al., 2021b; Wang et al., 2021e). We extend these findings here to show that the majority of antigen-specific B cells bind both WA1 and B.1.617.2 proteins. It is noteworthy that in NHP there is a more substantial contraction of the S-specific memory B cell compartment following vaccination than reported for humans (Goel et al., 2021b). Species-specific differences and the high percentage of memory B cells that bound SARS-CoV-2 S in our NHP immediately after vaccination may account for these observations. Memory B cell expansion after challenge likely provides a mechanism for the anamnestic antibody response detected in BAL. Although we detected this expansion in blood, it probably reflects a similar local response in lung lymphoid tissue (Poon et al., 2021). Importantly, the extent of anamnestic B cell activation may be dependent on the viral challenge dose and consequent virus replication. The increase in total IgG and measles-binding titers following challenge provide further evidence of a rapid polyclonal B cell response in the lung, likely driven by Toll-like receptor 7 (TLR7) activation by SARS-CoV-2, a single-stranded RNA virus (Bortolotti et al., 2021; Hornung et al., 2002). Indeed, transient innate stimulation of B cells in the lung may be critical for the initial adaptive immune response to respiratory pathogens and appears to precede T cell expansion.

There are multiple potential roles for T cells following vaccination by mRNA. This vaccine platform has been shown by us and others in NHP and humans to induce T_H_1, T_FH_ and a low frequency of CD8^+^ T cells. T_FH_ cells are important for inducing and sustaining antibody responses (Corbett et al., 2021b; Lederer et al., 2020; Painter et al., 2021; Pardi et al., 2018). Here we show that S-specific T cells decreased over the course of one year but were still detectable in a subset of vaccinated NHP at the time of challenge. A second potential role for T cells would be mediating control of viral replication in the lower airway to limit severe disease. Notably, we did not see increased T_H_1 or CD8^+^ T cell responses in the BAL of vaccinated NHP compared to controls following challenge. Interestingly, we detected CD8^+^ T cells in the blood before detection in the lungs post-challenge, which requires further investigation. Formal demonstration for whether T cells have a role in protection in the lungs would require depletion at the time of challenge.

A potential limitation relates to the extent of viral replication following B.1.617.2 challenge. The challenge stock used here was passaged once and fully matched the canonical B.1.617.2 S sequences obtained from humans. However, we did not observe greater sgRNA copy numbers in the upper airway than in previous challenges with B.1.351 or WA1 (Corbett et al., 2021b; Corbett et al., 2021c). Clinical reports have described substantially higher viral titers in the upper respiratory tract from B.1.617.2 infections compared to ancestral strains (Ong et al., 2021; Williams et al., 2021). It is possible that the NHP model does not precisely recapitulate human infection.

We and others have previously shown that antibodies are a correlate of protection (Corbett et al., 2021b; Gilbert et al., 2021). These studies are based on a short interval between vaccination and virus exposure or challenge. Here, we did not find a clear immune correlate of protection between binding or neutralizing antibody responses in the blood at the peak of the response or the time of challenge one year later. Additional analysis from human clinical trials with much larger numbers of vaccinated individuals than the 8 NHP in this study will be important for determining whether serum antibody titers remain as a correlate of protection in the long term. Indeed, the data presented here raise the possibility that the correlate of protection may lie in the ability of tissue-resident memory B cells to expand after infection.

In summary, mRNA-1273 provided durable protection against B.1.617.2 challenge in the lower airways likely through anamnestic induction of antibody responses in the lung. An important consideration is that control of virus was both limited in the nose and briefly delayed in the lungs, which together may provide the virus with greater opportunity for transmission, particularly if variants emerge that may be more transmissible, more virulent or more resistant to neutralization.

## METHOD DETAILS

### Cells and Viruses

VeroE6 cells were obtained from ATCC (clone E6, ATCC, #CRL-1586) and cultured in complete DMEM medium consisting of 1x DMEM (VWR, #45000-304), 10% FBS, 25mM HEPES Buffer (Corning Cellgro), 2mM L-glutamine, 1mM sodium pyruvate, 1x Non-essential Amino Acids, and 1x antibiotics. VeroE6-TMPRSS2 cells were generated at Vaccine Research Center, NIH, Bethesda, MD. Isolation and sequencing of EHC-083E (D614G SARS-CoV-2), B.1.351 and B.1.617.2 for live virus neutralization assays were previously described (Edara et al., 2021a; Edara et al., 2021b; Vanderheiden et al., 2021). Viruses were propagated in Vero-TMPRSS2 cells to generate viral stocks. Viral titers were determined by focus-forming assay on VeroE6-TMPRSS2 cells. Viral stocks were stored at −80°C until use.

### Sequencing of Virus Stock

We used NEBNext Ultra II RNA Prep reagents (New England Biolabs) to prepare Illumina-ready libraries, which were sequenced on a NextSeq 2000 (Illumina) as described previously (Corbett et al., 2021c). Demultiplexed sequence reads were analyzed in the CLC Genomics Workbench v.21.0.3 by (1) trimming for quality, length, and adaptor sequence, (2) mapping to the Wuhan-Hu-1 SARS-CoV-2 reference (GenBank accession number: NC_045512), (3) improving the mapping by local realignment in areas containing insertions and deletions (indels), and (4) generating both a sample consensus sequence and a list of variants. Default settings were used for all tools.

### Rhesus Macaque Model, Immunizations, and Delta Challenge

All experiments conducted according to NIH regulations and standards on the humane care and use of laboratory animals as well as the Animal Care and Use Committees of the NIH Vaccine Research Center and Bioqual, Inc. (Rockville, Maryland). 4- to 14-year-old Indian-origin rhesus macaques were housed at Bioqual, Inc. NHP were stratified based on age, weight, and gender into 2 cohorts (8 NHP per group). One group received 100µg mRNA-1273 at weeks 0 and 4, while the other group received 100µg of an mRNA control at weeks 0 and 4, as described previously (Corbett et al., 2021a). mRNA-1273 and mRNA controls were prepared in 1mL PBS. Vaccinations were given intramuscularly (IM) in the right quadriceps. 49 weeks after prime, NHP were challenge with 2×10^5^ PFU of SARS-CoV-2 B.1.617.2. 1.5×10^5^ PFU was resuspended in 3mL PBS and administered intratracheally, while 0.5×10^5^ PFU was resuspended in 1mL PBS and administered intranasally, with half of the volume in each nostril.

### Serum and Mucosal Antibody Titers

Determination of antibody responses in the blood and mucosa were performed as previously described for measurement of mucosal antibody responses (Corbett et al., 2020). Briefly, heat inactivated plasma was serially diluted 1:4. BAL fluid and nasal washes were concentrated 10-fold with Amicon Ultra centrifugal filter device (Millipore Sigma) and then serially diluted 1:5. Total IgG and IgA antigen-specific antibodies to variant SARS-CoV-2 RBD-derived antigens were determined by enzyme-linked immunosorbent assay (ELISA) using MULTI-ARRAY 384-well streptavidin-coated plates (Meso Scale Discovery, MSD). Plates were coated with variant-specific biotinylated antigens (see Table S1) at a concentration of 0.18μg/mL. MSD plate reader (Sector Imager 600) used for measurement of chemiluminescence. WHO international units were calculated using the MSD Reference Standard 1 against the WHO International Standard (NIBSC code: 20/136).

Total IgG antibody titers in the BAL were quantitated by using the Human/NHP IgG Kit (MSD) and antibody titers to measles were quantitated by using the Monkey Anti-Measles IgG ELISA Kit (Alpha Diagnostics International) according to the manufacturer’s instructions.

### S-2P-ACE2 Binding Inhibition

Heat inactivated plasma was diluted 1:40. BAL fluid and nasal washes were concentrated 10-fold with Amicon Ultra centrifugal filter devices (Millipore Sigma) and then diluted 1:5. ACE2 binding inhibition assay was performed with V-Plex SARS-CoV-2 Panel 13 (ACE2) Kit (MSD) per manufacturer’s instructions.

### Focus Reduction Neutralization Assay

FRNT assays were performed as previously described (Vanderheiden et al., 2020). Briefly, samples were diluted at 3-fold in 8 serial dilutions using DMEM (VWR, #45000-304) in duplicates with an initial dilution of 1:10 in a total volume of 60μl. Serially diluted samples were incubated with an equal volume of SARS-CoV-2 (100-200 foci per well) at 37°C for 1 hour in a round-bottomed 96-well culture plate. The antibody-virus mixture was then added to Vero cells and incubated at 37°C for 1 hour. Post-incubation, the antibody-virus mixture was removed and 100µl of prewarmed 0.85% methylcellulose (Sigma-Aldrich, #M0512-250G) overlay was added to each well. Plates were incubated at 37°C for 24 hours. After 24 hours, methylcellulose overlay was removed, and cells were washed three times with PBS. Cells were then fixed with 2% paraformaldehyde in PBS (Electron Microscopy Sciences) for 30 minutes. Following fixation, plates were washed twice with PBS and 100µl of permeabilization buffer (0.1% BSA [VWR, #0332], Saponin [Sigma, 47036-250G-F] in PBS), was added to the fixed Vero cells for 20 minutes. Cells were incubated with an anti-SARS-CoV S primary antibody directly conjugated to biotin (CR3022-biotin) for 1 hour at room temperature. Next, the cells were washed three times in PBS and avidin-HRP was added for 1 hour at room temperature followed by three washes in PBS. Foci were visualized using TrueBlue HRP substrate (KPL, # 5510-0050) and imaged on an ELISPOT reader (CTL).

Antibody neutralization was quantified by counting the number of foci for each sample using the Viridot program (Katzelnick et al., 2018). The neutralization titers were calculated as follows: 1 - (ratio of the mean number of foci in the presence of sera and foci at the highest dilution of respective sera sample). Each specimen was tested in duplicate. The FRNT_50_ titers were interpolated using a 4-parameter nonlinear regression in GraphPad Prism 8.4.3. Samples that do not neutralize at the limit of detection at 50% are plotted at 15, which was used for geometric mean calculations.

### Lentiviral Pseudovirus Neutralization

Pseudotyped lentiviral reporter viruses were produced as previously described (Wu et al., 2021). Briefly, HEK293T/17 cells (ATCC CRL-11268) were transfected with the following: plasmids encoding S proteins from Wuhan-Hu-1 strain (GenBank no. MN908947.3) with a p.Asp614Gly mutation, a luciferase reporter, lentivirus backbone, and the human transmembrane protease serine 2 (TMPRSS2) genes. For pseudovirus encoding the S from B.1.351 and B.1.617.2, the plasmid was altered via site-directed mutagenesis to match the S sequence to the corresponding variant sequence as previously described (Corbett et al., 2021a). Plasma and/or sera, in duplicate, were tested for neutralizing activity against the pseudoviruses by quantification of luciferase activity in relative light units (RLU). The percentage of neutralization was normalized, with luciferase activity in uninfected cells defined as 100% neutralization and luciferase activity in cells infected with pseudovirus alone as 0% neutralization. IC_50_ titers were calculated using a log(agonist) versus normalized-response (variable slope) nonlinear regression model in Prism v9.0.2 (GraphPad). For samples that did not neutralize at the limit of detection at 50%, a value of 20 was plotted and used for geometric mean calculations.

### Serum Antibody Avidity

Avidity was measured as described previously (Francica et al., 2021) in an adapted ELISA assay. Briefly, ELISA against S-2P was performed in the absence or presence of sodium thiocyanate (NaSCN), and the avidity index (AI) was calculated by determining the ratio of IgG binding to S-2P in the absence or presence of NaSCN. The reported AI is the average of two independent experiments, each containing duplicate samples.

### Epitope Mapping

Serum epitope mapping competition assays were performed using the Biacore 8K+ surface plasmon resonance system (Cytiva) as previously described (Corbett et al., 2021a). Briefly, anti-histidine antibody was immobilized on Series S Sensor Chip CM5 (Cytiva) through primary amine coupling using a His capture kit (Cytiva), allowing His-tagged SARS-CoV-2 S protein containing 2 proline stabilization mutations (S-2P) to be captured on active sensor surface.

Competitor human IgG mAbs used for these analyses include: RBD-specific mAbs B1-182, CB6, A20-29.1, LY-COV555, A19-61.1, and S309. Negative control antibody or competitor monoclonal antibody (mAb) was injected over both active and reference surfaces. Then, NHP sera (diluted 1:100) was flowed over both active and reference sensor surfaces. Active and reference sensor surfaces were regenerated between each analysis cycle.

Beginning at the serum association phase, sensorgrams were aligned to Y (Response Units) = 0, using Biacore 8K Insights Evaluation Software (Cytiva). Relative “analyte binding late” report points (RU) were collected and used to calculate relative percent competition (% C) using the following formula: % C = [1 – (100 * ((RU in presence of competitor mAb) / (RU in presence of negative control mAb)))]. Results are reported as percent competition. Assays were performed in duplicate, with average data point represented on corresponding graphs.

### B Cell Probe Binding

Previously isolated and cryopreserved PBMC were centrifuged into warm RPMI/10% FBS and resuspended in wash buffer (4% heat inactivated newborn calf serum/0.02% NaN_3_/ phenol-free RPMI). PBMC were transferred to 96-wells, washed twice in 1X PBS, and incubated with Aqua Blue live/dead cell stain (Thermo Fisher Scientific) at room temperature for 20 minutes. Following washing, cells were incubated with primary antibodies for 20 minutes at room temperature. The following antibodies were used (monoclonal unless indicated): IgD FITC (goat polyclonal, Southern Biotech), IgM PerCP-Cy5.5 (clone G20-127, BD Biosciences), IgA Dylight 405 (goat polyclonal, Jackson Immunoresearch Inc), CD20 BV570 (clone 2H7, Biolegend), CD27 BV650 (clone O323, Biolegend), CD14 BV785 (clone M5E2, Biolegend), CD16 BUV496 (clone 3G8, BD Biosciences), IgG Alexa 700 (clone G18-145, BD Biosciences), CD3 APC-Cy7 (clone SP34.2, BD Biosciences), CD38 PE (clone OKT10, Caprico Biotechnologies), CD21 PE-Cy5 (clone B-ly4, BD Biosciences), and CXCR5 PE-Cy7 (clone MU5UBEE, Thermo Fisher Scientific). Cells were washed twice in wash buffer and then incubated with streptavidin-BV605 (BD Biosciences) labeled B.1.617.2 S-2P and streptavidin-BUV661 (BD Biosciences) labeled WA1 S-2P for 30 minutes at 4°C (protected from light). Cells were washed twice in wash buffer and residual RBC were lysed using BD FACS Lysing Solution (BD Biosciences) at room temperature for 10 minutes. Following two final washes, cells were fixed in 0.5% formaldehyde (Tousimis Research Corp). All antibodies titrated to determine the optimal concentration. Samples were acquired on an BD FACSymphony cytometer and analyzed using FlowJo version 10.7.2 (BD, Ashland, OR).

### Intracellular Cytokine Staining

Cryopreserved PBMC and BAL cells were thawed and rested overnight in a 37°C/5% CO2 incubator. The next morning, cells were stimulated with SARS-CoV-2 S protein (S1 and S2, matched to vaccine insert) and N peptide pools (JPT Peptides) at a final concentration of 2 μg/ml in the presence of 3mM monensin for 6 hours. The S1, S2 and N peptide pools are comprised of 158, 157 and 102 individual peptides, respectively, as 15mers overlapping by 11 aa in 100% DMSO. Negative controls received an equal concentration of DMSO instead of peptides (final concentration of 0.5%). Intracellular cytokine staining was performed as described (Donaldson et al., 2019). The following monoclonal antibodies were used: CD3 APC-Cy7 (clone SP34.2, BD Biosciences), CD4 PE-Cy5.5 (clone S3.5, Invitrogen), CD8 BV570 (clone RPA-T8, BioLegend), CD45RA PE-Cy5 (clone 5H9, BD Biosciences), CCR7 BV650 (clone G043H7, BioLegend), CXCR5 PE (clone MU5UBEE, Thermo Fisher), CXCR3 BV711 (clone 1C6/CXCR3, BD Biosciences), PD-1 BUV737 (clone EH12.1, BD Biosciences), ICOS Pe-Cy7 (clone C398.4A, BioLegend), CD69 ECD (cloneTP1.55.3, Beckman Coulter), IFN-g Ax700 (clone B27, BioLegend), IL-2 BV750 (clone MQ1-17H12, BD Biosciences), IL-4 BB700 (clone MP4-25D2, BD Biosciences), TNF-FITC (clone Mab11, BD Biosciences), IL-13 BV421 (clone JES10-5A2, BD Biosciences), IL-17 BV605 (clone BL168, BioLegend), IL-21 Ax647 (clone 3A3-N2.1, BD Biosciences), and CD154 BV785 (clone 24-31, BioLegend). Aqua live/dead fixable dead cell stain kit (Thermo Fisher Scientific) was used to exclude dead cells. All antibodies were previously titrated to determine the optimal concentration. Samples were acquired on a BD FACSymphony flow cytometer and analyzed using FlowJo version 10.8.0 (Treestar, Inc., Ashland, OR).

BAL samples from weeks 6 and 25 were stimulated with S1 peptide pools only. One BAL sample from a vaccinated NHP on Day 2 post-challenge had extensive background staining for CD8 markers and was excluded from analysis of both S and N responses.

### Titration of Delta Stock in Syrian Hamsters

All experiments conducted according to NIH regulations and standards on the humane care and use of laboratory animals as well as the Animal Care and Use Committees of the NIH Vaccine Research Center and Bioqual, Inc. (Rockville, Maryland). 6-week-old male Syrian hamsters (Envigo) were housed at Bioqual, Inc. Hamsters were stratified based on weight into 3 cohorts of 4 hamsters each. Hamsters were infected with B.1.617.2 challenge doses of 10^5^, 10^4^, and 10^3^ PFU diluted in 100µL PBS and split between both nostrils. Weight changes and clinical observations were collected daily.

### Titration of Delta Stock in Nonhuman Primates

All experiments conducted according to NIH regulations and standards on the humane care and use of laboratory animals as well as the Animal Care and Use Committees of the NIH Vaccine Research Center and Bioqual, Inc. (Rockville, Maryland). 4-year-old male Indian-origin rhesus macaques were housed at Bioqual, Inc. NHP were stratified based on weight into 2 cohorts (3 NHP per group). Groups received 1×10^5^ or 1×10^6^ PFU of SARS-CoV-2 B.1.617.2. 75% of the challenge dose was resuspended in 3mL PBS and administered intratracheally, while the remaining virus was resuspended in 1mL PBS and administered intranasally, with half of the volume in each nostril. BAL fluid and nasal swabs were collected on days 2, 5 and 7 following challenge for sgRNA quantification.

### Subgenomic RNA Quantification

sgRNA was isolated and quantified as previously described (Corbett et al., 2021c). Briefly, total RNA was extracted from BAL fluid and nasal swabs using RNAzol BD column kit (Molecular Research Center). PCR reactions were conducted with TaqMan Fast Virus 1-Step Master Mix (Applied Biosystems), forward primer in the 5’ leader region, and the following gene-specific probes and reverse primers.

sgLeadSARSCoV2_F: 5’-CGATCTCTTGTAGATCTGTTCTC-3’

*E gene*

E_Sarbeco_P: 5’-FAM-ACACTAGCCATCCTTACTGCGCTTCG-BHQ1-3’

E_Sarbeco_R: 5’-ATATTGCAGCAGTACGCACACA-3’

*N gene*

wtN_P: 5’-FAM-TAACCAGAATGGAGAACGCAGTGGG-BHQ1-3’

wtN_R: 5’-GGTGAACCAAGACGCAGTAT-3’

Amplifications were performed with a QuantStudio 6 Pro Real-Time PCR System (Applied Biosystems). The assay lower limit of detection was 50 copies per reaction.

### TCID_50_ Assay

Vero-TMPRSS2 cells (obtained from Adrian Creanga, Vaccine Research Center-NIAID) were plated at 25,000 cells/well in DMEM with 10% FBS and gentamicin, and the cultures were incubated at 37°C, 5.0% CO_2_. Media was aspirated and replaced with 180μL of DMEM with 2% FBS and gentamicin. 10-fold serial dilutions of samples starting from 20μL of material were added to the cells in quadruplicate and incubated at 37°C for 4 days. Positive (virus stock of known infectious titer) and negative (medium only) controls were included in each assay. The plates were incubated at 37°C, 5.0% CO_2_ for 4 days. Cell monolayers were visually inspected for cytopathic effect (CPE). The TCID_50_ was calculated using the Read-Muench formula.

### Histopathology and Immunohistochemistry

Histopathology and immunohistochemistry (IHC) were performed as described previously (Corbett et al., 2020). Briefly, H&E staining and IHC were conducted and analyzed by a board-certified veterinary pathologist with an Olympus BX51 light microscope. A rabbit polyclonal anti-SARS-CoV-2 antibody (GeneTex, GTX135357) at a dilution of 1:2000 was used for IHC. Photomicrographs were taken on an Olympus DP73 camera.

### Statistical Analysis

NS denotes that the indicated comparison was not significant, with *P*>0.05. We did not perform statistical analysis at any timepoint in which we had samples from fewer than 5 NHP.

Comparisons between groups, or between timepoints within a group, are based on unpaired and paired t-tests, respectively. All analysis for serum epitope mapping was performed using unpaired, two-tailed t-test. Binding, neutralizing and viral assays are log-transformed as appropriate and reported with geometric means and corresponding confidence intervals where indicated. Correlations are based on Spearman’s nonparametric rho, and the associated asymptotic p-values. There are no adjustments for multiple comparisons, so all p-values and significance testing should be interpreted as suggestive rather than conclusive. All analyses are conducted using R version 4.0.2 and GraphPad Prism version 8.2.0 unless otherwise specified.

## RESOURCES AVAILABILITY

### Lead contact

Further information and requests for resources should be directed to and will be fulfilled by the lead contacts, Daniel C. Douek and Robert A. Seder.

### Materials availability

This study did not generate new unique reagents.

### Data and code availability

All data is in the main text or supplementary figures.

## ACKNOWLEDGEMENTS

We would like to thank G. Alvarado for experimental organization and administrative support. I.-T. Teng provided variant-specific antigens used for MULTI-ARRAY ELISA. The VRC Production Program (VPP) provided the WA1 protein for the avidity assay. We would also like to especially thank A. Olia and C. Liu for providing the B.1.617.2 protein and J. Misasi, C. Stringham, C. Schramm, Y. Zhang, M. Choe, A. Henry, and B. Zhang for providing critical reagents, sequencing analysis, or expert technical assistance for serum antibody epitope mapping assays. D. Flebbe, S. Provost, E. Lamb, J. Marquez, A. Mychalowych and M. Donaldson processed samples and prepared reagents for both ICS and B cell probe binding assays. A. Van Ry, B. Narvaez, Z. Flinchbaugh, and D. Valentin provided technical assistance for the TCID_50_ assays. We would like to thank M. Boursiquot and K. Steingrebe for assistance with NHP experiments. C. Case coordinated hamster experiments. M. Porto, C. Kitajewski, R. Stone, N. Daham, M. Hamilton, J. Wear, A. Faudree, B. Chang, J. Vazquez, and T. Wheatley provided technical assistance on hamster experiments. VPP contributors include C. Anderson, V. Bhagat, J. Burd, J. Cai, K. Carlton, W. Chuenchor, N. Clbelli, G. Dobrescu, M. Figur, J. Gall, H. Geng, D. Gowetski, K. Gulla, L. Hogan, V. Ivleva, S. Khayat, P. Lei, Y. Li, I. Loukinov, M. Mai, S. Nugent, M. Pratt, E. Reilly, E. Rosales-Zavala, E. Scheideman, A. Shaddeau, A. Thomas, S. Upadhyay, K. Vickery, A. Vinitsky, C. Wang, C. Webber, and Y. Yang.

This project has been funded in part by both the Intramural Program of the National Institute of Allergy and Infectious Diseases, National Institutes of Health, Department of Health and Human Services and under HHSN272201400004C (NIAID Centers of Excellence for Influenza Research and Surveillance, CEIRS) and NIH P51 OD011132 awarded to Emory University. This work was also supported in part by the Emory Executive Vice President for Health Affairs Synergy Fund award, COVID-Catalyst-I3 Funds from the Woodruff Health Sciences Center and Emory School of Medicine, the Pediatric Research Alliance Center for Childhood Infections and Vaccines and Children’s Healthcare of Atlanta, and Woodruff Health Sciences Center 2020 COVID-19 CURE Award.

## Author Contributions

M.R., M.C.N., N.J.S., D.C.D. and R.A.S. designed experiments. M.G., K.S.C, B.J.F., K.E.F., D.A.W., S.F.A., J-P.M.T., C.C.H., L.M., S.T.N., M.E.D-G., L.P., K.W.B., B.M.N., M.M., A.P.W., J.I.M., C.T., C.G.L., B.Z., E.M., A.C., A.D., P.M., J.R-T., F.L., L.W., A.G., S.K., A.P., N.D-R., A.C., J.R.M., E.A.B., D.K.E., H.A., M.G.L., M.S.S., B.S.G., M.R., I.N.M., M.C.N., N.J.S., D.C.D, and R.A.S. performed, analyzed, and/or supervised experiments. M.G., K.E.F., D.A.W., S.F.A., B.Z., E.A.B. and I.N.M. designed figures. L.W., S.B-B., E.S.Y., W.S., provided critical reagents. M.G., D.C.D. and R.A.S. wrote manuscript. All authors edited the manuscript and provided feedback on research.

