## Supplemental Material for "Protection from SARS-CoV-2 Delta one year after mRNA-1273 vaccination in nonhuman primates is coincident with an anamnestic antibody response in the lower airway"

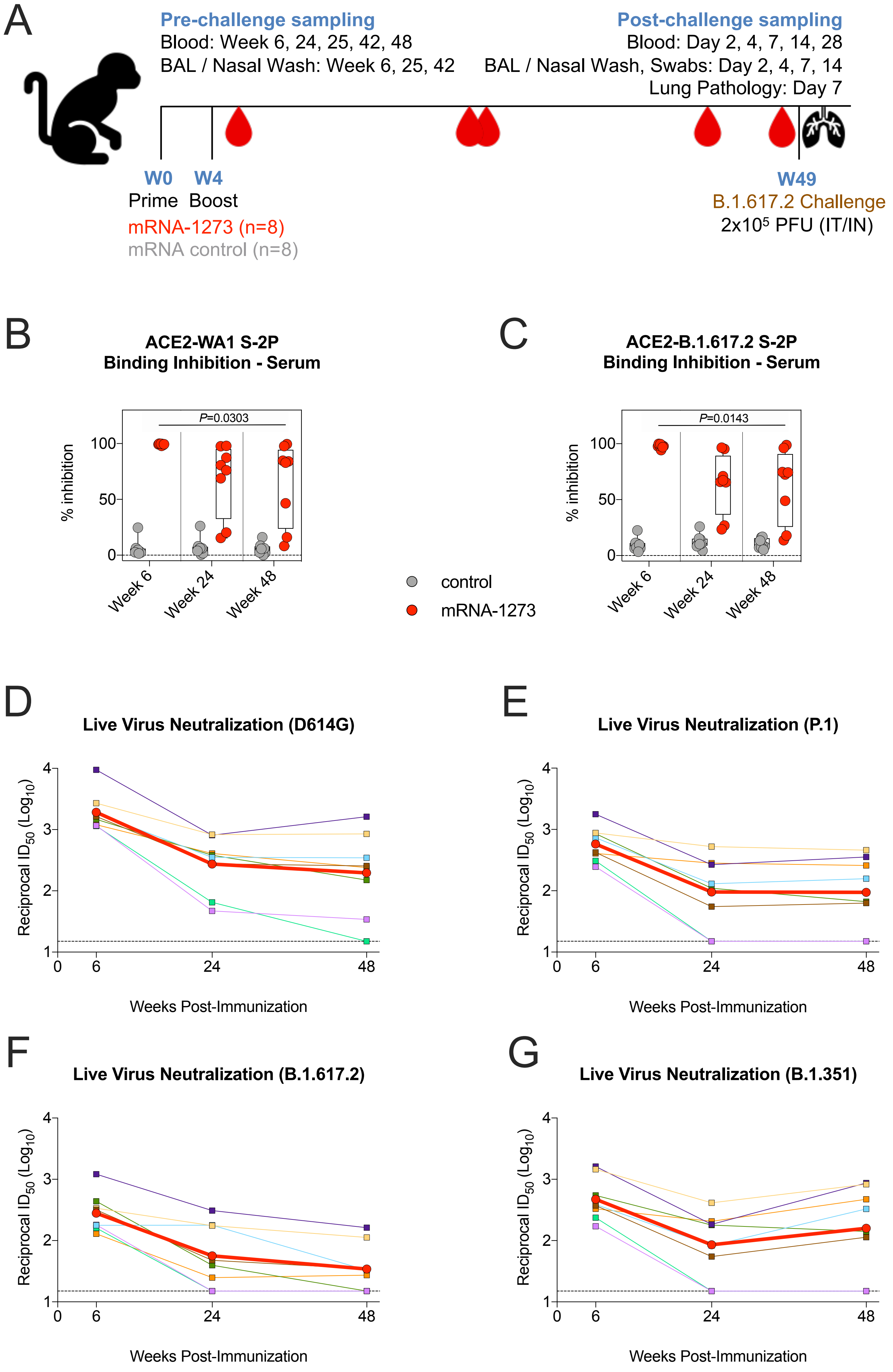
A
Pre-challenge sampling
Blood: Week 6, 24, 25, 42, 48
BAL / Nasal Wash: Week 6, 25, 42
Post-challenge sampling
Blood: Day 2, 4, 7, 14, 28
BAL / Nasal Wash, Swabs: Day 2, 4, 7, 14
Lung Pathology: Day 7
W0
Prime
W4
Boost
mRNA-1273 (n=8)
mRNA control (n=8)
W49
B.1.617.2 Challenge
2x10⁵ PFU (IT/IN)
B
ACE2-WA1 S-2P
Binding Inhibition - Serum
P=0.0303
% inhibition
100
50
0
Week 6
Week 24
Week 48
C
ACE2-B.1.617.2 S-2P
Binding Inhibition - Serum
P=0.0143
% inhibition
control
mRNA-1273
D
Live Virus Neutralization (D614G)
Reciprocal ID50 (Log10)
Weeks Post-Immunization
0
6
24
48
E
Live Virus Neutralization (P.1)
F
Live Virus Neutralization (B.1.617.2)
G
Live Virus Neutralization (B.1.351)

Figure S1

**Figure S1 Serum antibody kinetics after mRNA-1273 immunization**

(A) Experimental timeline showing immunization schedule and B.1.617.2 challenge. 8 NHP vaccinated with mRNA-1273, and 8 NHP given mRNA control.

(B-C) Sera were collected at weeks 6, 24 and 48 post-immunization and diluted 1:40. SARS-CoV-2 WA1 (B) and B.1.617.2 (C) S-2P binding to ACE2 measured both alone and in the presence of sera to calculate % inhibition. Circles denote individual NHP. Boxes represent interquartile range with the median denoted by a horizontal line. Dotted lines set to 0% inhibition. 8 NHP per group. Statistical analysis shown for mRNA-1273 cohort only.

(D-G) Sera were collected at weeks 6, 24 and 48 post-immunization. Reciprocal $ID_{50}$ titers calculated for neutralization of live virus D614G (D), P.1 (E), B.1.617.2 (F) and B.1.351 (G). Color-coded squares and lines indicate individual NHP. Red circles and thick lines indicate geometric means. Dotted lines indicate assay limit of detection. 8 NHP per group.

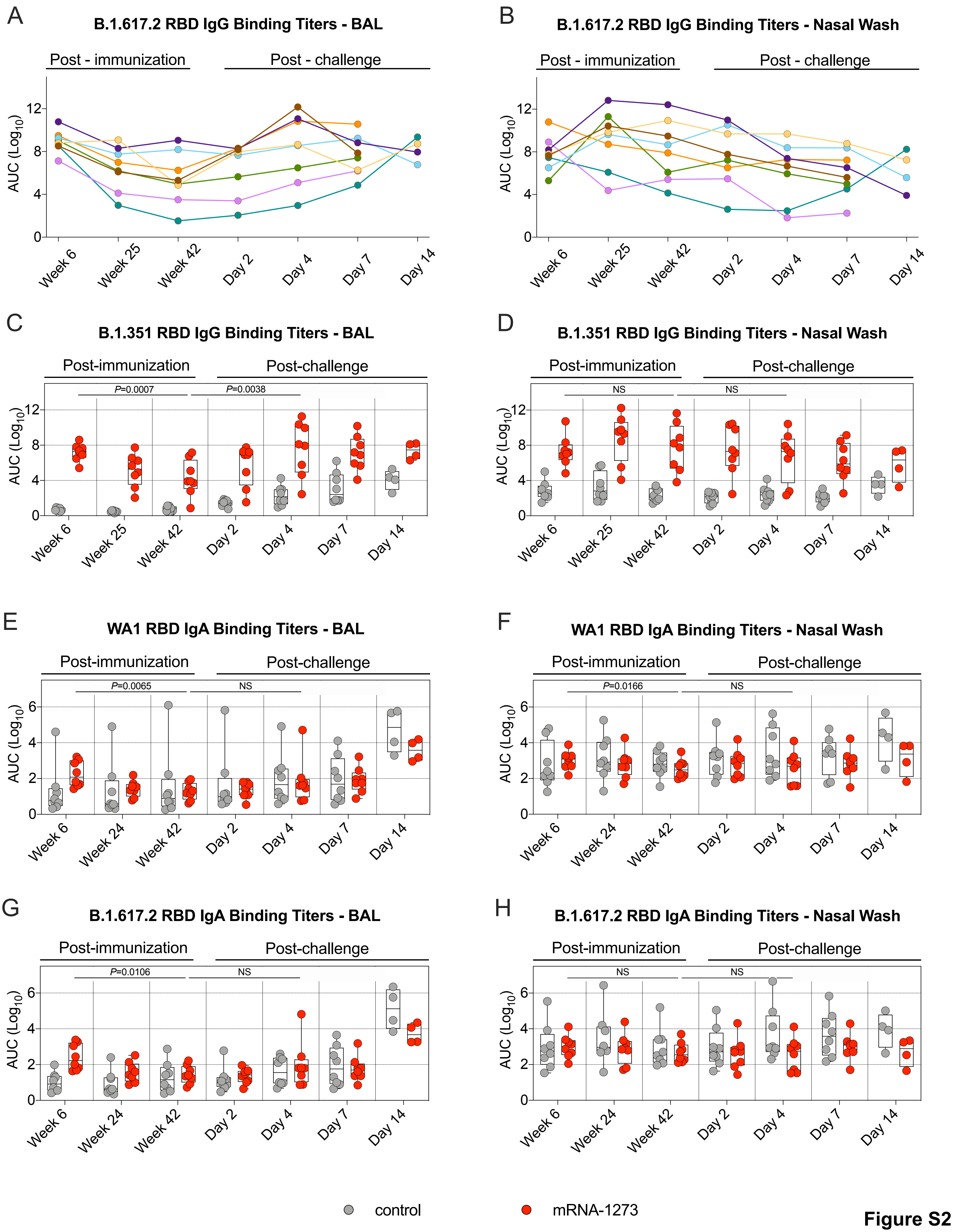

A
B.1.617.2 RBD IgG Binding Titers - BAL
Post - immunization
Post - challenge
AUC (Log10)
Week 6
Week 25
Week 42
Day 2
Day 4
Day 7
Day 14
B
B.1.617.2 RBD IgG Binding Titers - Nasal Wash
C
B.1.351 RBD IgG Binding Titers - BAL
Post-immunization
Post-challenge
P=0.0007
P=0.0038
D
B.1.351 RBD IgG Binding Titers - Nasal Wash
NS
E
WA1 RBD IgA Binding Titers - BAL
P=0.0065
Week 24
F
WA1 RBD IgA Binding Titers - Nasal Wash
P=0.0166
G
B.1.617.2 RBD IgA Binding Titers - BAL
P=0.0106
H
B.1.617.2 RBD IgA Binding Titers - Nasal Wash
control
mRNA-1273


Figure S2

**Figure S2 Kinetics of RBD-binding IgG and IgA responses in the upper and lower airway**

(A-D) BAL and nasal washes were collected at weeks 6, 25 and 42 post-immunization, and days 2, 4, 7 and 14 post-challenge. B.1.617.2 (A-B) and B.1.351 (C-D) RBD-binding IgG titers in the lower (A, C) or upper airway (B, D). Circles indicate individual NHP. 4-8 NHP per group. (A-B) Color-coded lines indicate temporal changes in B.1.617.2 RBD-binding titers for individual NHP. All animals received mRNA-1273. (C-D) Boxes represent interquartile range with the median denoted by a horizontal line. Dotted lines are for visualization purposes and denote 4-$\log_{10}$ increases in binding titers.

(E-H) BAL and nasal washes were collected at weeks 6, 24 and 42 post-immunization, and days 2, 4, 7 and 14 post-challenge. (E-F) WA1 and (G-H) B.1.617.2 RBD-binding IgA titers in the lower (E, G) or upper airway (F, H). Circles in (E-H) indicate individual NHP. Boxes represent interquartile range with the median denoted by a horizontal line. Dotted lines are for visualization purposes and denote 2-$\log_{10}$ increases in binding titers. 4-8 NHP per group.

Statistical analysis in (C-H) shown for mRNA-1273 cohort only.

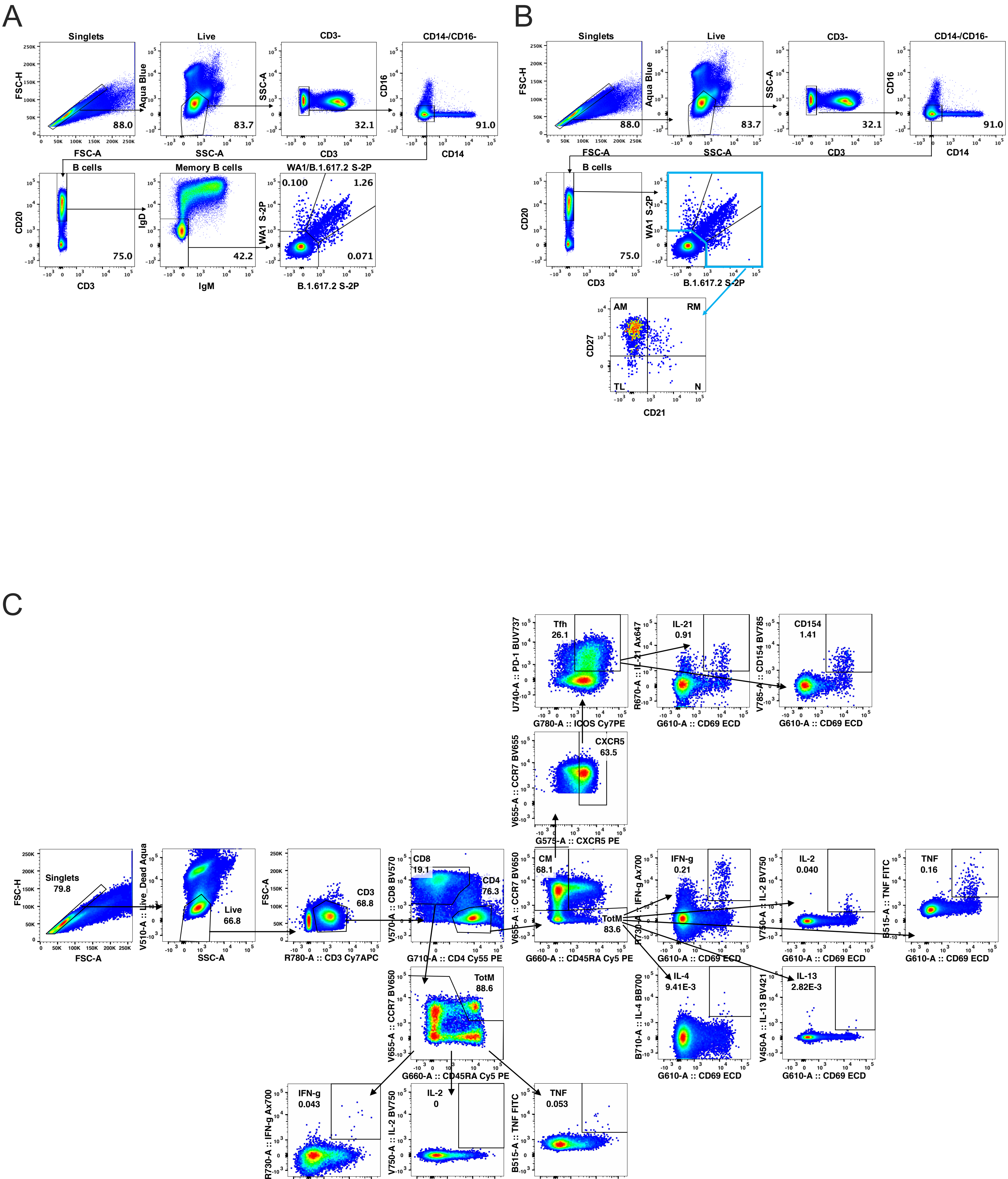
A
Singlets
FSC-H
FSC-A
88.0
Live
Aqua Blue
SSC-A
83.7
CD3-
SSC-A
CD3
32.1
CD14-/CD16-
CD16
CD14
91.0
B cells
CD20
CD3
75.0
Memory B cells
IgD
IgM
42.2
WA1/B.1.617.2 S-2P
WA1 S-2P
B.1.617.2 S-2P
0.100
1.26
0.071
B
Singlets
FSC-H
FSC-A
88.0
Live
Aqua Blue
SSC-A
83.7
CD3-
SSC-A
CD3
32.1
CD14-/CD16-
CD16
CD14
91.0
B cells
CD20
CD3
75.0
WA1 S-2P
B.1.617.2 S-2P
AM
RM
TL
N
CD27
CD21
C
Singlets
79.8
FSC-H
FSC-A
Live
66.8
V510-A :: Live_Dead Aqua
SSC-A
CD3
68.8
FSC-A
R780-A :: CD3 Cy7APC
CD8
19.1
CD4
76.3
V570-A :: CD8 BV570
G710-A :: CD4 Cy55 PE
CM
68.1
TotM
83.6
V655-A :: CCR7 BV650
G660-A :: CD45RA Cy5 PE
CXCR5
63.5
V655-A :: CCR7 BV655
G575-A :: CXCR5 PE
Tfh
26.1
U740-A :: PD-1 BUV737
G780-A :: ICOS Cy7PE
IL-21
0.91
R670-A :: IL-21 Ax647
G610-A :: CD69 ECD
CD154
1.41
V785-A :: CD154 BV785
G610-A :: CD69 ECD
IFN-g
0.21
R730-A :: IFN-g Ax700
G610-A :: CD69 ECD
IL-2
0.040
V750-A :: IL-2 BV750
G610-A :: CD69 ECD
TNF
0.16
B515-A :: TNF FITC
G610-A :: CD69 ECD
IL-4
9.41E-3
B710-A :: IL-4 BB700
G610-A :: CD69 ECD
IL-13
2.82E-3
V450-A :: IL-13 BV421
G610-A :: CD69 ECD
TotM
88.6
V655-A :: CCR7 BV650
G660-A :: CD45RA Cy5 PE
IFN-g
0.043
R730-A :: IFN-g Ax700
G610-A :: CD69 ECD
IL-2
0
V750-A :: IL-2 BV750
G610-A :: CD69 ECD
TNF
0.053
B515-A :: TNF FITC
610-A :: CD69 ECD

**Figure S3**

**Figure S3 B and T cell gating strategy**

(A) Representative flow cytometry plots showing gating strategy for B cells in Figure 4. Cells were gated as singlets and live cells on forward and side scatter and a live/dead aqua blue stain. CD3- cells were then gated on absence of CD14 and CD16 expression and positive CD20 expression. Memory B cells were selected based on lack of IgD or IgM. Finally, WA1 S-2P and B.1.617.2 S-2P probes were used to determine binding specificity.

(B) Representative flow cytometry plots showing gating strategy for B cell phenotypes in Figure 4F. $CD20^+$ B cells were gated as described above. WA1 S-2P and B.1.617.2 S-2P probes were used to determine binding specificity. Probe-binding cells were further characterized as having a phenotype consistent with naïve (N), tissue-like memory (TL), activated memory (AM), or resting memory (RM) cells according to expression of CD27 and CD21.

(C) Representative flow cytometry plots showing gating strategy for T cells in Figure 5 and Figure S4. Cells were gated as singlets and live cells on forward and side scatter and a live/dead aqua blue stain. $CD3^+$ events were gated as $CD4^+$ or $CD8^+$ T cells. Total memory $CD8^+$ T cells were selected based on expression of CCR7 and CD45RA. Finally, SARS-CoV-2 S-specific memory $CD8^+$ T cells were gated according to co-expression of CD69 and IL-2, TNF or IFNγ. The $CD4^+$ events were defined as naïve, total memory or central memory according to expression of CCR7 and CD45RA. $CD4^+$ cells with a $T_H1$ phenotype were defined as memory cells that co-expressed CD69 and IL-2, TNF or IFNγ. $CD4^+$ cells with a $T_H2$ phenotype were defined as memory cells that co-expressed CD69 and IL-4 or IL-13. $T_{FH}$ cells were defined as central memory $CD4^+$ T cells that coexpressed CXCR5, ICOS and PD-1. $T_{FH}$ cells were further characterized as IL-$21^+$, $CD69^+$ or $CD40L^+$, $CD69^+$.

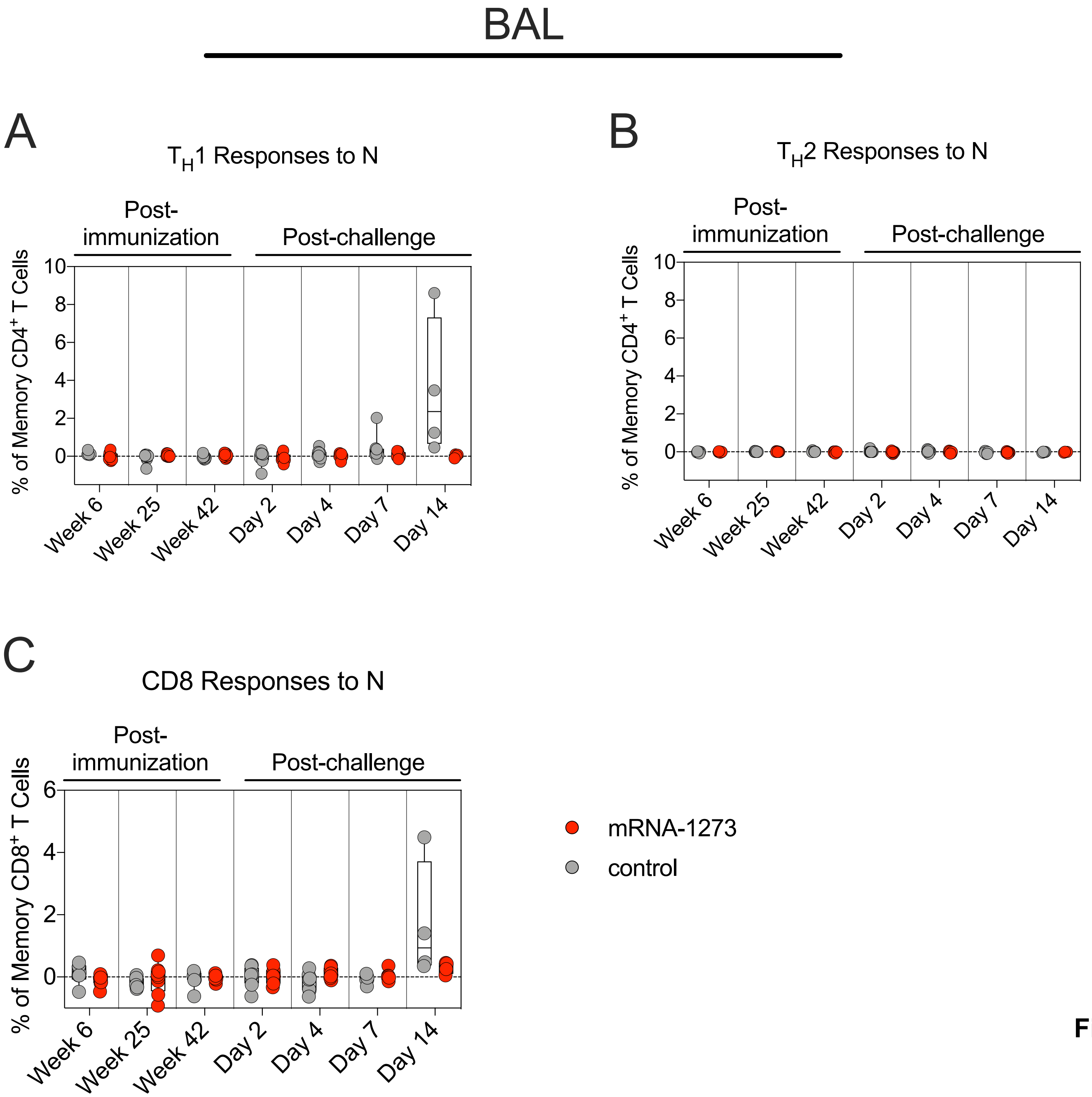


**Figure S4**

**Figure S4 Vaccinated NHP do not mount a primary T cell response to SARS-CoV-2 N peptides following challenge**

(A-C) BAL fluid was collected at weeks 6, 25 and 42 post-immunization, and days 2, 4, 7 and 14 post-challenge. Lymphocytes in the BAL were stimulated with N peptide pools and responses measured by intracellular cytokine staining. (A-B) Percentage of memory $CD4^+$ T cells with (A) $T_H1$ markers (IL-2, TNF or IFNγ) or (B) $T_H2$ markers (IL-4 or IL-13) following stimulation. (C) Percentage of $CD8^+$ T cells expressing IL-2, TNF or IFNγ. Circles in (A-C) indicate individual NHP. Boxes represent interquartile range with the median denoted by a horizontal line. Dotted lines set at 0%. Reported percentages may be negative due to background subtraction. 4-8 NHP per group. See also Figure S3 for gating strategy.

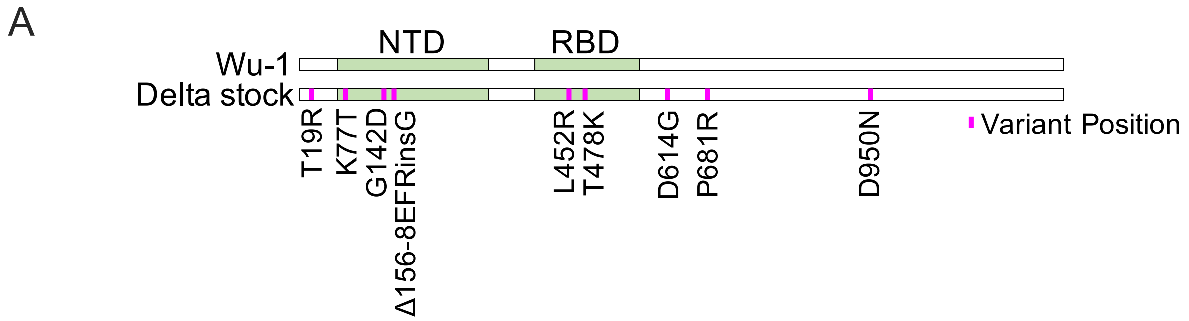


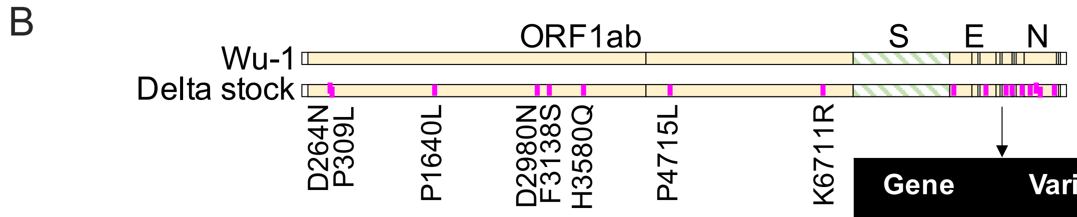


| Gene | Variant |
|---|---|
| ORF3a | S26L |
| M | I82T |
| ORF7a | P45FS* (Frameshift) |
| ORF7a | T120I |
| ORF8 | C102F |
| N | D63G |
| N | P142Q |
| N | R203M |
| N | D377Y |

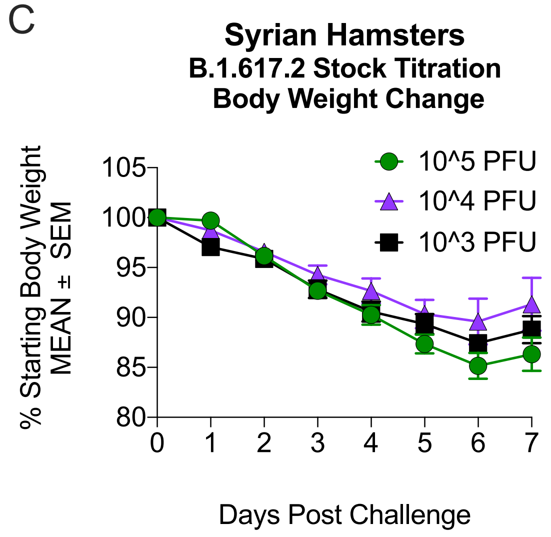


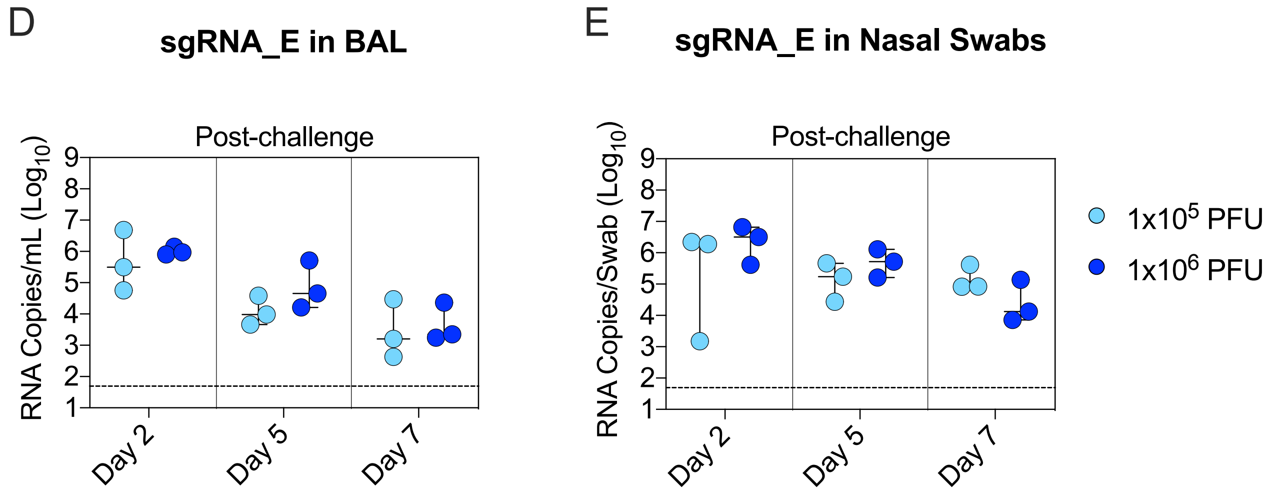


**Figure S5**

**Figure S5 B.1.617.2 challenge stock characterization**

(A-B) Alignment of B.1.617.2 stock to Wuhan-Hu-1 reference sequence. (A) S gene only. (B) Whole genome. Variant positions depicted in purple. Multiple mutations at the 3' end of genome shown in table for visualization purposes. *Non-canonical frameshift in ORF7a produced by 172 nucleotide deletion. Deletion matched original isolate.

(C) B.1.617.2 stock titration in golden Syrian hamsters at 3 doses spanning 2-logs PFU. Arithmetic mean body weight across indicated days in comparison to starting weight shown. Error bars indicate standard error of the mean. 4 hamsters per group.

(D-E) B.1.617.2 stock titration in rhesus macaques at dose of $1x10^5$ or $1x10^6$ PFU. Copies sgRNA_E per mL BAL (D) or per NS (E). Circles indicate individual NHP. Median value denoted by horizontal line. Dotted lines indicate assay limit of detection. 3 NHP per group.

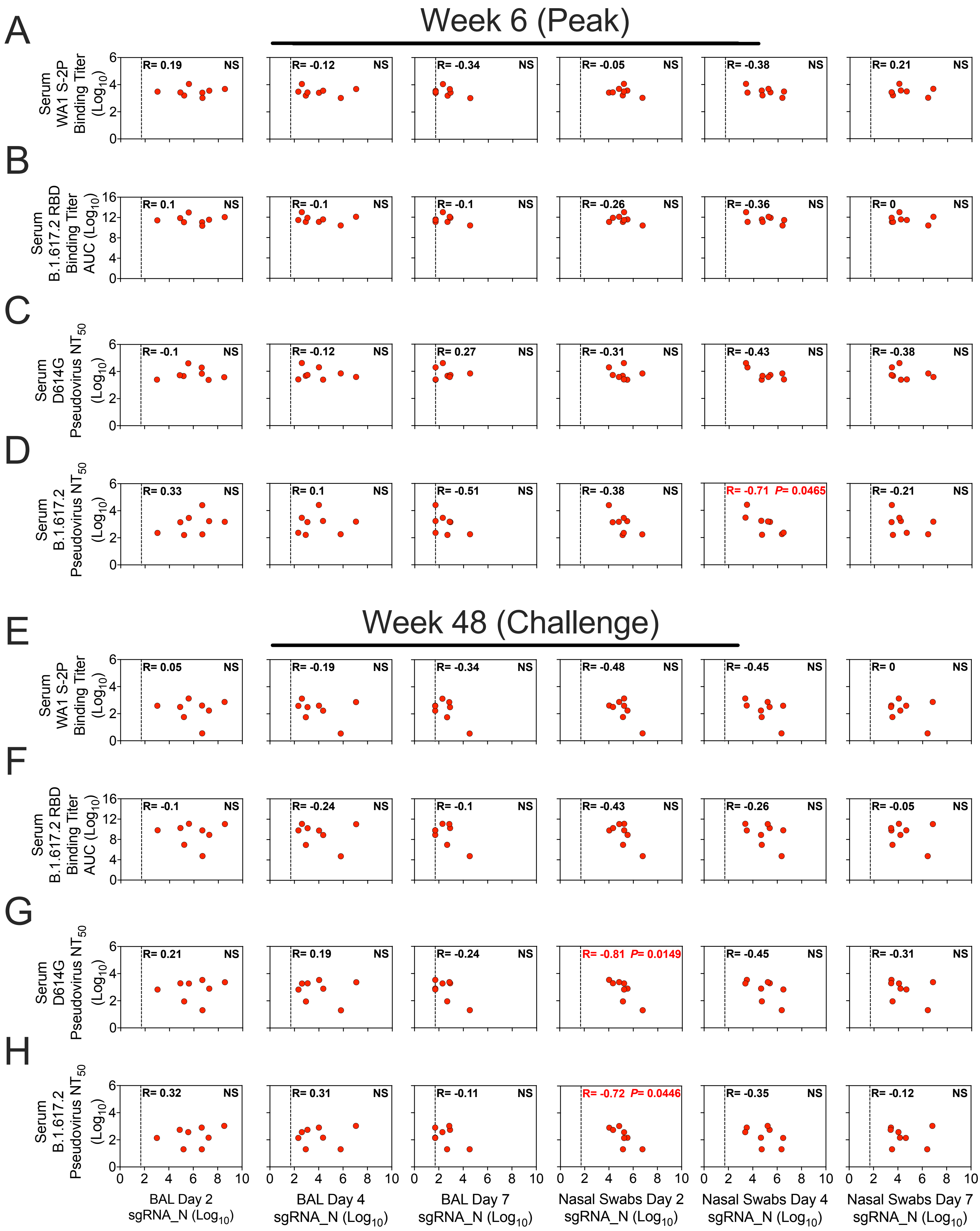
A
Week 6 (Peak)
Serum WA1 S-2P Binding Titer (Log10)
R= 0.19 NS
R= -0.12 NS
R= -0.34 NS
R= -0.05 NS
R= -0.38 NS
R= 0.21 NS
B
Serum B.1.617.2 RBD Binding Titer AUC (Log10)
R= 0.1 NS
R= -0.1 NS
R= -0.1 NS
R= -0.26 NS
R= -0.36 NS
R= 0 NS
C
Serum D614G Pseudovirus NT50 (Log10)
R= -0.1 NS
R= -0.12 NS
R= 0.27 NS
R= -0.31 NS
R= -0.43 NS
R= -0.38 NS
D
Serum B.1.617.2 Pseudovirus NT50 (Log10)
R= 0.33 NS
R= 0.1 NS
R= -0.51 NS
R= -0.38 NS
R= -0.71 P= 0.0465
R= -0.21 NS
E
Week 48 (Challenge)
Serum WA1 S-2P Binding Titer (Log10)
R= 0.05 NS
R= -0.19 NS
R= -0.34 NS
R= -0.48 NS
R= -0.45 NS
R= 0 NS
F
Serum B.1.617.2 RBD Binding Titer AUC (Log10)
R= -0.1 NS
R= -0.24 NS
R= -0.1 NS
R= -0.43 NS
R= -0.26 NS
R= -0.05 NS
G
Serum D614G Pseudovirus NT50 (Log10)
R= 0.21 NS
R= 0.19 NS
R= -0.24 NS
R= -0.81 P= 0.0149
R= -0.45 NS
R= -0.31 NS
H
Serum B.1.617.2 Pseudovirus NT50 (Log10)
R= 0.32 NS
R= 0.31 NS
R= -0.11 NS
R= -0.72 P= 0.0446
R= -0.35 NS
R= -0.12 NS
BAL Day 2 sgRNA_N (Log10)
BAL Day 4 sgRNA_N (Log10)
BAL Day 7 sgRNA_N (Log10)
Nasal Swabs Day 2 sgRNA_N (Log10)
Nasal Swabs Day 4 sgRNA_N (Log10)
Nasal Swabs Day 7 sgRNA_N (Log10)

**Figure S6**

**Figure S6 Serum antibody titers are not a correlate for protection one year after vaccination**

(A-H) Correlations between sgRNA_N copies per mL of BAL or per NS at days 2, 4 and 7 post-challenge and indicated binding or neutralizing titers. Antibody responses include serum binding titers to WA1 S-2P in WHO units (A, E), serum binding titers to B.1.617.2 RBD (B, F), serum lentiviral pseudovirus neutralizing titers to D614G (C, G), or serum lentiviral neutralizing titers to B.1.617.2 (D, H) at week 6 (A-D) or week 48 (E-H) post-immunization. R denotes Spearman correlation coefficient. Dotted line indicates qRT-PCR limit of detection. Correlates analysis limited to vaccinated NHP.

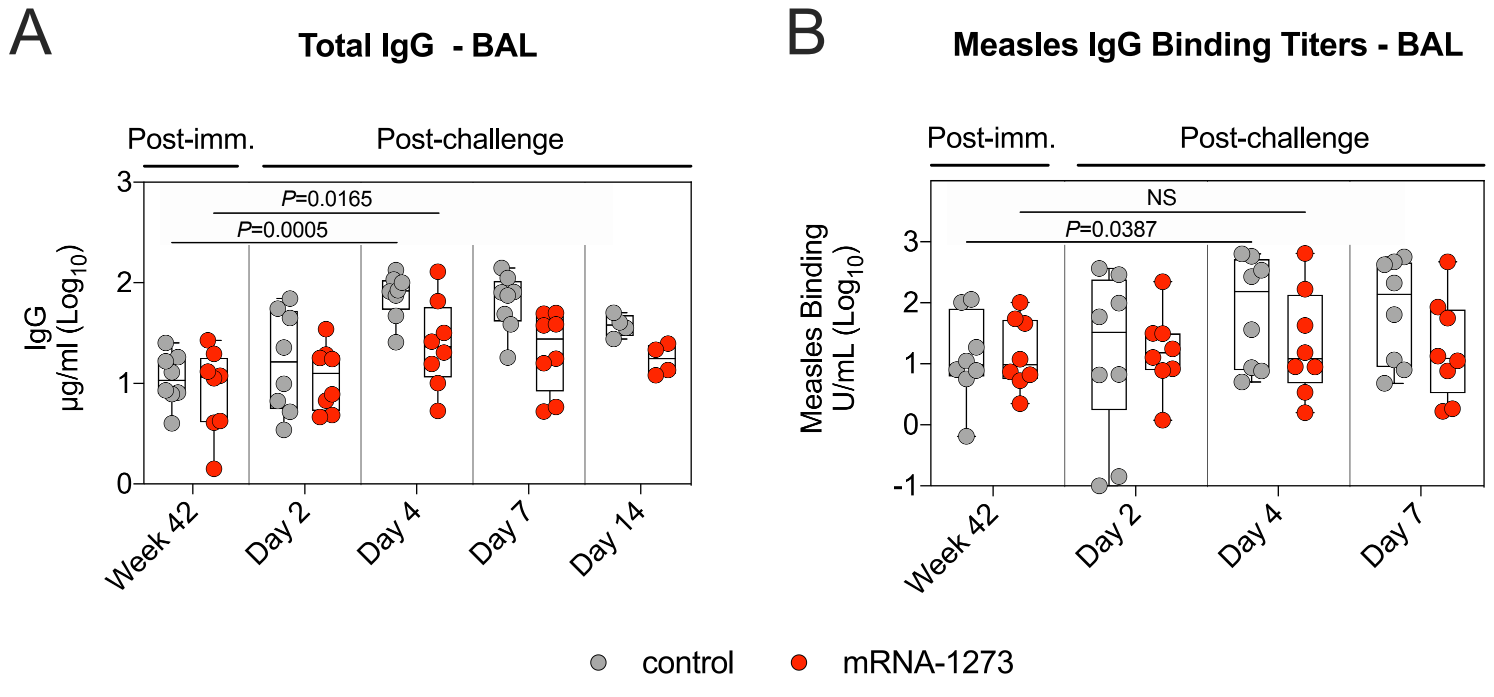


**Figure S7**

**Figure S7 B.1.617.2 challenge induces a broad antiviral immune response**

(A-B) BAL fluid was collected at week 42 post-immunization, and days 2, 4, 7 and 14 post-challenge. (A) Total IgG. (B) Measles morbillivirus-binding titers. Circles in (A-B) indicate individual NHP. Boxes represent interquartile range with the median denoted by a horizontal line. 4-8 NHP per group. Statistical analysis shown for values at week 42 in comparison to day 4 post-challenge for both control NHP (lower line) and mRNA-1273 cohort (upper line).

| Variant | Mutations in RBD |
| --- | --- |
| **WA1** | None |
| **B.1.351** | K417N, E484K, N501Y |
| **B.1.617.2** | L452R, T478K |

**Table S1 List of mutations in variant-specific MULTI-ARRAY ELISA**

All variant positions in reference to Wuhan-Hu-1 sequence (Genbank: MN908947.3).

| **Subdomain** | **Site** | **mAb** | |
|---|---|---|---|
| **RBD** | | | **Barnes classification (Barnes et al., 2020)** |
| | **A** | B1-182 + | CLASS I |
| | **B** | CB6 + | |
| | **C** | A20-29.1 | CLASS II |
| | **E** | LY-COV555 | |
| | **F** | A19-61.1 + | CLASS III |
| | **G** | S309 | |

**Table S2 SARS-CoV2 S Antigenic Sites**

Monoclonal antibodies (mAbs) used to define cross-reactive antigenic sites on SARS-CoV-2 S are defined by non-competing binding on SARS-CoV-2 WA1 S. RBD-targeted mAbs have been grouped based on their Barnes classification (Barnes et al., 2020). Monoclonal antibodies which have been shown to be neutralizing to B.1.617.2 variant of SARS-CoV-2 are denoted with "+".

| | | | sgRNA_N (Copies / mL or Copies / swab) | | | | | |
|---|---|---|---|---|---|---|---|---|
| Time Point | Site | Assay | Day 2 BAL | Day 4 BAL | Day 7 BAL | Day 2 NS | Day 4 NS | Day 7 NS |
| Week 6 | Serum | WA1 S-2P Titers (BAU/mL) | 0.19 | -0.12 | -0.34 | -0.05 | -0.38 | 0.21 |
| Week 48 | Serum | WA1 S-2P Titers (BAU/mL) | 0.05 | -0.19 | -0.34 | -0.48 | -0.45 | 0 |
| Week 6 | Serum | B.1.617.2 RBD IgG Titers (AUC) | 0.1 | -0.1 | -0.1 | -0.26 | -0.36 | 0 |
| Week 48 | Serum | B.1.617.2 RBD IgG Titers (AUC) | -0.1 | -0.24 | -0.1 | -0.43 | -0.26 | -0.05 |
| Week 6 | BAL | B.1.617.2 RBD IgG Titers (AUC) | 0.17 | 0.02 | -0.17 | -0.38 | -0.67 | -0.4 |
| Week 42 | BAL | B.1.617.2 RBD IgG Titers (AUC) | -0.4 | -0.52 | -0.37 | -0.45 | -0.43 | -0.57 |
| Week 6 | Nasal Wash | B.1.617.2 RBD IgG Titers (AUC) | -0.1 | -0.24 | -0.22 | -0.52 | -0.45 | -0.21 |
| Week 42 | Nasal Wash | B.1.617.2 RBD IgG Titers (AUC) | -0.12 | -0.31 | -0.22 | -0.33 | -0.21 | 0.07 |
| Week 6 | BAL | B.1.617.2 RBD IgA Titers (AUC) | 0.29 | 0.1 | -0.39 | -0.02 | -0.07 | 0.38 |
| Week 42 | BAL | B.1.617.2 RBD IgA Titers (AUC) | -0.26 | -0.24 | -0.05 | 0.12 | 0.14 | -0.17 |
| Week 6 | Nasal Wash | B.1.617.2 RBD IgA Titers (AUC) | -0.21 | 0 | -0.1 | -0.55 | 0.36 | -0.02 |
| Week 42 | Nasal Wash | B.1.617.2 RBD IgA Titers (AUC) | -0.1 | 0.12 | 0.17 | -0.64 | 0.19 | -0.14 |
| Week 6 | Serum | D614G Pseudovirus NT ($ID_{50}$) | -0.1 | -0.12 | 0.27 | -0.31 | -0.43 | -0.38 |
| Week 48 | Serum | D614G Pseudovirus NT ($ID_{50}$) | 0.21 | 0.19 | -0.24 | **-0.81** | -0.45 | -0.31 |
| Week 6 | Serum | B.1.617.2 Pseudovirus NT ($ID_{50}$) | 0.33 | 0.1 | -0.51 | -0.38 | **-0.71** | -0.21 |
| Week 48 | Serum | B.1.617.2 Pseudovirus NT ($ID_{50}$) | 0.32 | 0.31 | -0.11 | **-0.72** | -0.35 | -0.12 |

**Table S3 Correlates analyses of antibody responses and virus subgenomic copy numbers**

Correlations between antibody responses in the blood, BAL, and nasal washes at week 6 (peak) or at week 42-48 (memory and time of challenge) to sgRNA_N copy numbers in the BAL and nose at days 2, 4, and 7 after B.1.617.2 challenge. Values listed are Spearman correlation coefficient (R). R values in red denote correlations that are statistically significant ($P<0.05$). All analyses are exploratory, and significant associations should be viewed as suggestive.
